# A cGAMP-containing hydrogel for prolonged SARS-CoV-2 RBD subunit vaccine exposure induces a broad and potent humoral response

**DOI:** 10.1101/2021.07.03.451025

**Authors:** Volker Böhnert, Emily C. Gale, Lauren J. Lahey, Jerry Yan, Abigail E. Powell, Ben S. Ou, Jacqueline A. Carozza, Lingyin Li, Eric A. Appel

## Abstract

The receptor binding domain (RBD) of the SARS-CoV-2 virus spike protein has emerged as a promising target for generation of neutralizing antibodies. Although the RBD subunit is more stable than its encoding mRNA, RBD-based subunit vaccines have been hampered by RBD’s poor immunogenicity. We hypothesize that this limitation can be overcome by sustained co-administration with a more potent and optimized adjuvant than standard adjuvants. The endogenous innate immune second messenger, cGAMP, holds promise as potent activator of the anti-viral STING pathway. Unfortunately, delivery of cGAMP as a therapeutic exhibits poor performance due to poor pharmacokinetics and pharmacodynamics from rapid excretion and degradation by its hydrolase ENPP1. To overcome these limitations, we sought to create an artificial immunological niche enabling slow release of cGAMP and RBD to mimic natural infections in which immune activating molecules are co-localized with antigen. Specifically, we co-encapsulated cGAMP and RBD in an injectable polymer-nanoparticle (PNP) hydrogel. This cGAMP-adjuvanted hydrogel vaccine elicited more potent, durable, and broad antibody responses with improved neutralization as compared to dose-matched bolus controls and hydrogel-based vaccines lacking cGAMP. The cGAMP-adjuvanted hydrogel platform developed is suitable for delivery of other antigens and may provide enhanced immunity against a broad range of pathogens.

## 1. Introduction

Subunit vaccines with recombinant protein antigens are a safe and scalable approach for preventing infectious disease.^[1]^ In the context of a pandemic, such as the COVID-19 pandemic, notable advantages of subunit vaccines include their favorable safety profile, high stability at warmer temperatures, and ease of manufacture at existing facilities across the globe.^[2]^ ^[3]^ ^[4]^ ^[5]^ With respect to SARS-CoV-2, the receptor-binding domain (RBD) of the spike protein on the viral surface is a critical epitope required for engaging host cells to initiate infection and is therefore a promising candidate antigen. RBD is easily and efficiently produced (up to 100-fold higher expression compared to spike trimer), is relatively stable, and is the target for 90% of serum neutralizing activity in humans.^[2]^ ^[6]^ ^[7]^ ^[8]^ ^[9]^ Unfortunately, RBD is poorly immunogenic and must be delivered with one or more adjuvant(s) to elicit an effective immune response. Moreover, while there has been progress towards drug delivery solutions for subunit vaccines to focus on improving the spatiotemporal delivery of vaccine components to the immune system, very few platforms exist that enable co-delivery of diverse antigens and adjuvant molecules, even though both sustained co-delivery and the inclusion of potent adjuvants are known to boost immune responses.^[10]^ ^[11]^ There is a critical need for development of potent adjuvant systems affording spatiotemporal control over vaccine exposure to enhance immune responses to poorly immunogenic subunit antigens such as the SARS-CoV-2 RBD protein.

2’3’-cyclic-GMP-AMP (cGAMP) is a molecular second messenger that acts as a potent agonist of the stimulator of interferon genes (STING) receptor resulting in upregulation of type I interferons (IFN) and defense responses.^[12]^ Endogenous cGAMP production can result from diverse cellular threats, such as viral infection and cancer, and plays a powerful role in activating innate immunity.^[13]^ Several innate and adaptive immune cells have been described to either directly or indirectly respond to cGAMP, including subsets of NK cells, macrophages, and dendritic cells (DCs), ultimately boosting elicitation of downstream adaptive immune responses to neutralize the cellular threat.^[14]^ ^[15]^ ^[16]^ ^[17]^ Given the roles of endogenous cGAMP, using exogenous cGAMP as a therapeutic represents a promising strategy to adjuvant vaccines and cancer immunotherapies. However, cGAMP, as a second messenger, is rapidly degraded by its hydrolase ENPP1, which is prevalent in the plasma.^[14]^ ^[15]^ To overcome its poor stability, intratumoral injection of a non-hydrolyzable cGAMP analog, ADU-S100, entered clinical trials (NCT03172936, NCT03937141) to evaluate its ability to increase tumor immunogenicity and synergize with checkpoint inhibitors. However, this non-hydrolyzable cGAMP analog did not deliver satisfying clinical results partially because of its poor pharmacokinetics and pharmacodynamics. At low injection concentrations, it rapidly diffuses from the site of administration causing a lack of efficacy at the desired tumor site.^[18]^ While injection of a high concentration of a non-degradable cGAMP analog represents one strategy to temporarily elevate the concentration at the injection site, this approach is associated with dose-dependent systemic toxicity, and adverse effects.^[19]^ Employing non-hydrolyzable cGAMP analogs as effective and safe vaccine adjuvants is even more challenging due to the lack of an immunological niche equivalent to the tumor microenvironment, in addition to their poor pharmacokinetics and pharmacodynamics.^[20]^ ^[21]^We propose to harness natural cGAMP as a safe and effective adjuvant using a hydrogel platform that localizes and sustains low but steady cGAMP and antigen concentrations and also serves as an immunological niche; any cGAMP leaked out of the niche should be rapidly degraded by ENPP1 to minimize systemic interferon responses.

Many efforts to control cGAMP delivery have focused on engineering particle-based systems that passively drain to lymph nodes where they are taken up by cells. pH-responsive polymerosomes,^[22]^ acetylated dextran microparticles,^[23]^ polymer nanoparticles (e.g. polyethylenimine/hyaluronic acid; poly(beta-amino ester),^[24]^ ^[25]^ and engineered pulmonary surfactant-biomimetic liposomes^[26]^ are all methods that have been pursued with differing success. While these cGAMP delivery systems have demonstrated promise, each still has important limitations including low encapsulation efficiency, complex manufacturing, and relatively poor stability. Most of these approaches induce endosomal uptake of cGAMP-loaded particles, which has the potential for non-specific uptake and could lead to STING activation in cell types associated with unwanted toxicity. We and others recently showed that different cell types have different cGAMP importer profiles and therefore different abilities to take up free extracellular cGAMP.^[27]^ ^[28]^ ^[29]^ ^[30]^ Differences in cell-type specific cGAMP uptake efficiency may help explain the efficacy of viral infection-induced cGAMP production at which concentrations only selective and desirable cellular cGAMP import occurs. We envision that a successful cGAMP delivery platform mimicking the natural slow release of soluble cGAMP would be highly effective, simple to manufacture at large scale, and easily loaded with diverse antigen cargo for utility across numerous disease indications.^[5]^

To overcome the limitations of current subunit vaccine delivery systems while achieving slow-release of soluble cGAMP, our lab developed an injectable polymer-nanoparticle (PNP) hydrogel that is simple to make, scalable, and readily loaded with diverse cargo. These hydrogel vaccines act through two main mechanisms: (i) vaccine cargo are released slowly over timescales relevant to a natural viral infection and (ii) immune cells infiltrate the hydrogel to interact with high local levels of both antigen and adjuvant in a de novo immune niche.^[31]^ ^[32]^ In this work, we evaluated whether our PNP hydrogel vaccine delivery platform could overcome the delivery challenges of cGAMP, improving cGAMP’s capacity to act as an adjuvant in the context of a SARS-CoV-2 RBD subunit vaccine. We envisioned the hydrogel system could mirror biological contexts such as viral infection or cancer in which endogenous cGAMP production is localized, sustained, and acts as a powerful immune potentiator.^[16]^ ^[33]^ Herein, we show that a subcutaneous hydrogel immunization containing cGAMP, the commonly used adjuvant Alhydrogel (alum), and RBD achieves a potent and durable humoral response in mice. As compared to dose-matched bolus controls, hydrogel vaccines led to significantly higher anti-RBD antibody titers that were maintained against recent SARS-CoV-2 variants of concern. Antibody subtype titers demonstrated that inclusion of cGAMP in the hydrogel or bolus immunizations improved skewing towards a Th1 (*i.e.*, cell-mediated response), which is thought to be central to fighting the SARS-CoV-2 virus.^[34]^ Moreover, sera from hydrogel vaccinated animals possessed significantly greater neutralizing ability compared to bolus controls as evaluated by a pseudotyped SARS-CoV-2 infectivity assay. Finally, we demonstrated that inclusion of cGAMP as an adjuvant in the hydrogel improves recruitment of immune cells to the hydrogel vaccine niche to promote development of a rapid anti-RBD antibody response. Together, these results establish our PNP hydrogel as an effective delivery system for cGAMP in the context of an RBD-based SARS-CoV-2 subunit vaccine.

## 2. Results

### 2.1. Injectable PNP hydrogels enable sustained release of cGAMP

We have previously reported that PNP hydrogels, which are readily formed by mixing hydrophobically-modified hydroxypropylmethylcellulose (HPMC-C12) and biodegradable polymeric nanoparticles (NPs) made of poly(ethylene glycol)-*b*-poly(lactic acid), can enhance the immunogenicity and safety of model antigens and clinically de-risked adjuvants by controlling their delivery.^[31]^ ^[32]^ ^[35]^ ^[36]^ Here, we employed the PNP hydrogel system to encapsulate SARS-CoV-2 RBD antigen and cGAMP as an adjuvant (**Figure 1**a). We also chose to include alum, which has served as a gold-standard base adjuvant in hepatitis A and B, diptheria-tetanus, and pneumococcal vaccines, and is known to adsorb negatively charged proteins and molecules to further promote controlled release.^[37]^ We also included alum in all hydrogel groups because we were primarily interested in preserving vaccine identity (i.e. cGAMP, alum, and RBD) and investigating whether delivery from our hydrogel provided enhanced titers and neutralization compared to the dose-matched bolus equivalents. Although our focus is on improving cGAMP delivery, our broader goal is to develop a modular hydrogel delivery platform that is compatible with existing subunit vaccines, many of which include alum. The resulting hydrogel-based vaccines are easily injected using standard syringe and needle, and self-heal following extrusion through a needle to form a solid depot in the subcutaneous space (**Figure 1**b).^[36]^ ^[38]^ These hydrogels exhibit a unique ability to inhibit passive diffusion of entrapped cargo, thereby prolonging the cargo’s release, while permitting active motility of immune cells, enabling the materials to serve as an inflammatory niche into which immune cells traffic to encounter antigen and adjuvant at high local concentrations.^[5]^ ^[32]^ ^[39]^

**Figure 1.**
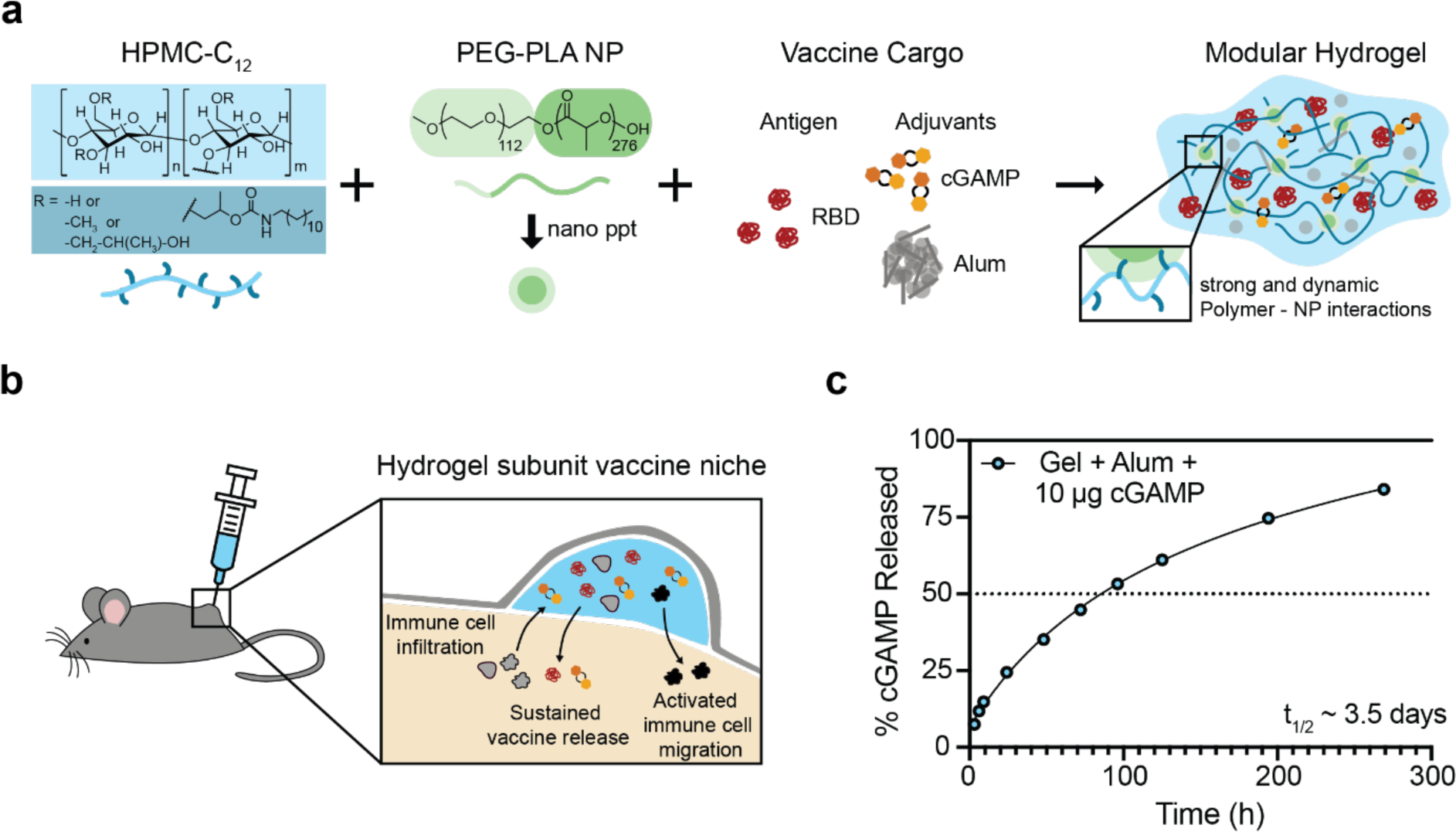
Polymer-nanoparticle (PNP) hydrogels provide prolonged co-delivery and generate an immune cell-activating niche for subunit RBD vaccine components. (A) Dodecyl-modified hydroxypropylmethylcellulose (HPMC-C12) is combined by simple mixing with poly(ethylene glycol)-*b*-poly(lactic acid) (PEG-PLA) and vaccine cargo (RBD, cGAMP, and alum) to form PNP hydrogels. Dynamic and multivalent noncovalent interactions between the HPMC-C12 polymer and NPs leads to reversible physical crosslinking within the hydrogel. (B) Hydrogel vaccines are injected subcutaneously, forming a solid-like depot under the skin that releases vaccine components slowly over time and provides an inflammatory niche for immune cell infiltration.^[2]^ ^[6]^ ^[9]^ ^[31]^ ^[32]^ The RBD protein was chosen as the antigen because it is stable, easy to manufacture, and is well-conserved in emerging SARS-CoV-2 variants.^[2]^ ^[6]^ ^[9]^ (C) Percent cGAMP released over time from a hydrogel vaccine in infinite sink conditions *in vitro* (n = 3 from one experiment). Mean +/- SD displayed along with “One site – Total” curve fit in GraphPad Prism.

We first assessed the *in vitro* release kinetics of vaccine components from the hydrogel. Our prototype vaccine hydrogels were loaded into capillary tubes and incubated with saline buffer at 37°C. The buffer sink was completely exchanged at the indicated times and the amount of vaccine component released into solution quantified in each sample. Controlled release of cGAMP from the hydrogel resulted in a release half-life of roughly 3.5 days (**Figure 1c**), which is similar to the half-life of RBD release *in vivo* previously observed (t1/2 = 5.5 days).^[31]^ Based on the diffusivity of cGAMP in PBS determined using the Stokes-Einstein equation, the half-life for cGAMP diffusion from the injection site when delivered in a PBS bolus can be estimated to be approximately 3 minutes, further supporting the hypothesis that the PNP hydrogel system can improve pharmacokinetics of cGAMP delivery by significantly slowing the rate of diffusion from the injection site.^[40]^

### 2.2. cGAMP-adjuvanted hydrogels promote acute IFN signaling followed by rapid and consistent elicitation of a durable anti-SARS-CoV-2 RBD antibody response

We next wanted to evaluate the effect of controlling cGAMP and RBD delivery from hydrogels on elicitation of humoral immunity. We therefore prepared vaccine variants in either bolus or PNP hydrogel forms that contained RBD (10 μg), alum (115 μg), and cGAMP (10 μg or 50 μg). Additionally, we sought to compare the adjuvant effect of hydrogel-delivered cGAMP versus that of poly(I:C) (50 μg), a high-molecular weight polymeric toll-like receptor 3 (TLR3) and RIG-I like receptor (RLR) agonist commonly employed in vaccines and evaluated in our original demonstration of PNP hydrogel-based vaccines.^[32]^ Vaccines were subcutaneously administered to 8-week old C57BL/6 mice at week 0 (prime) and week 8 (boost), with sera collected at weeks 1-12 (**Figure 2**a).

**Figure 2.**
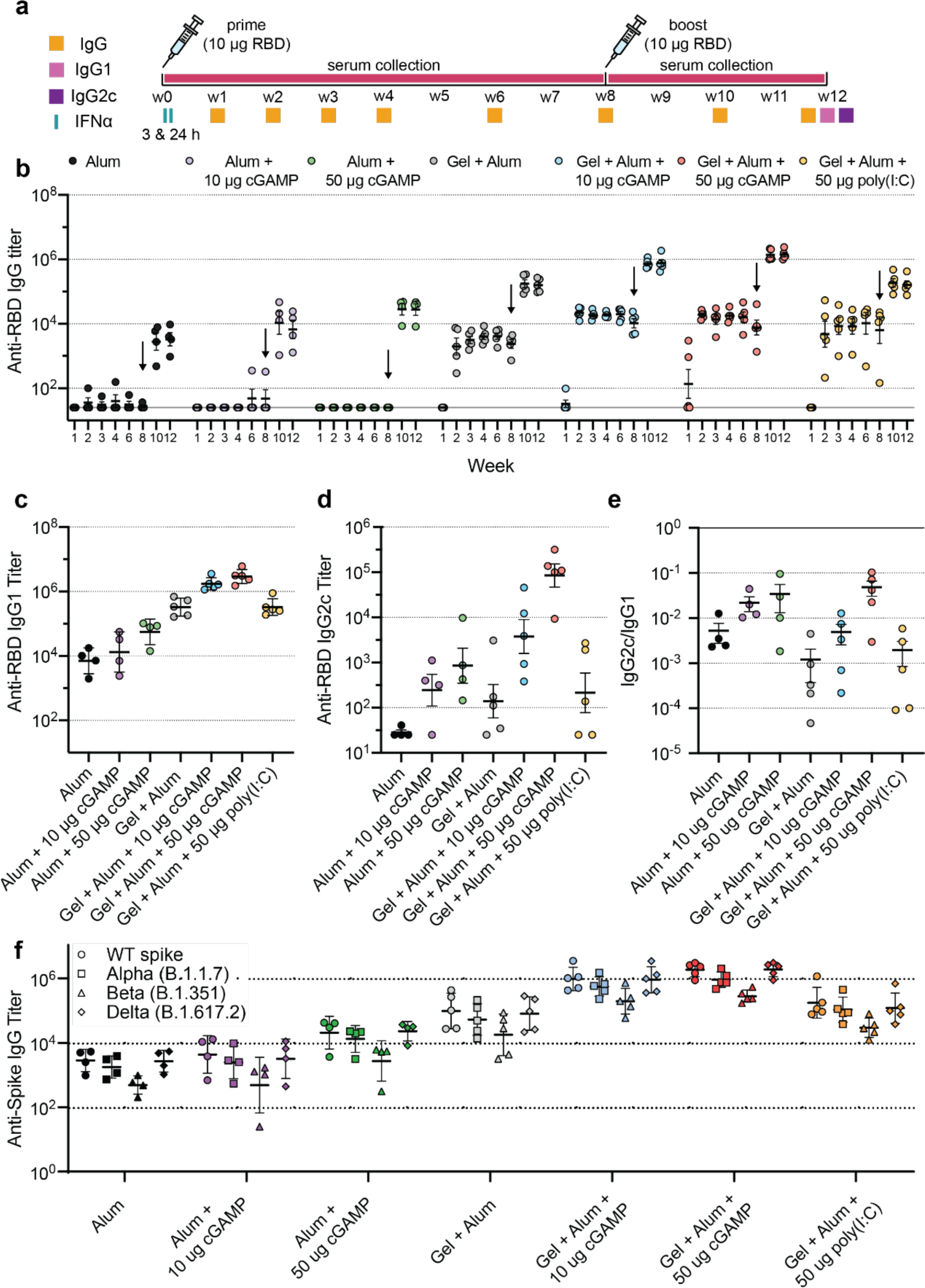
cGAMP-adjuvanted hydrogels elicit rapid, robust antibody responses against RBD and spike variants. (A) Vaccination schedule and timing of ELISAs. Mice were immunized at week 0 and week 8, serum was collected periodically over 12 weeks and assays were performed as indicated. (B) Weekly anti-RBD total IgG ELISA titers before and after boosting (arrow). (C-D) Anti-RBD IgG1 (C) and IgG2c (D) titers from post-boost, week 12 serum. (E) Calculated ratio of anti-RBD IgG2c to IgG1 post-boost titers. Values closer to 1 suggest a balanced Th2/Th1 response whereas values well-below 1 suggest Th2 skewing towards a stronger humoral response. (F) Post-boost anti-spike IgG titers against the native spike trimer as well as the Alpha (B.1.1.7), Beta (B.1.351), and Delta (B.1.617.2) variants. (B-F) Each point represents an individual mouse (n = 4-5). Data are shown as mean +/- SEM. P values listed in the text were determined using a 2way ANOVA with either Dunnett’s or Tukey’s multiple comparisons test (depending on number of groups compared) on the logged titer values for IgG titer comparisons (**Table S3-5)**. P values were determined using a 2way ANOVA with Dunnett’s multiple comparisons test on logged titer values for IgG1, IgG2c, and spike variant titer comparisons (**Table S6-7**).

cGAMP activation of STING leads to production of type I interferons consisting of the IFNα cytokine family, IFNβ, and others, which trigger expression of genes that upregulate the effector function of immune cells like dendritic cells (DCs), T cells and B cells.^[41]^ For this reason, we assessed the acute IFNα response in sera 3- and 24-hours after the initial immunization (**Figure S1**a, b, **Tables S1** and **S2**). We observed cGAMP dose-dependent IFNα levels in serum at 3- and 24-hours and negligible levels of IFNα in the absence of cGAMP. The 10 and 50 µg cGAMP hydrogel groups had slightly elevated levels of IFNα compared to the dose-matched bolus groups at the early 3-hour time point but had similar IFNα levels by 24 hours. Strikingly, IFNα levels following vaccination with poly(I:C), which is also known to be a potent type I interferon producer, were undetectable at 3h and low at 24h. We were not able to detect the pro-inflammatory cytokines CXCL10 and TNFα in any sample at 24h, indicating that cGAMP leads very specifically to systemically detectable type I IFN induction. Together, these results support the notion that cGAMP, and particularly cGAMP delivered in our hydrogel, serves as a potent IFNα-inducing adjuvant.

To analyze the kinetics of antibody development, we measured anti-RBD total IgG endpoint titers across eight timepoints (**Figure 2a**, b). With a single bolus vaccine dose, antibody titers were low or below the limit of detection for most mice, irrespective of cGAMP inclusion. In contrast, at two weeks following a control hydrogel prime dose, the average anti-RBD endpoint titer was 3.5×10^3^, with addition of poly(I:C) increasing the response by 4.4-fold (P = 0.34) and addition of 10 μg or 50 μg cGAMP increasing the response by 6.6-fold (P = 0.0004) or 5.6-fold (P = 0.001), respectively (**Table S3**). Notably, hydrogels including cGAMP as an adjuvant resulted in anti-RBD titers that rose most rapidly and had the smallest titer deviation across animals, both of which are desirable vaccine characteristics. The adjuvant effect of cGAMP in the hydrogel was sustained over the two months following a single vaccine dose: at weeks 4 and 8, cGAMP-hydrogel titers remained elevated, by 4.2-to 4.7-fold over the control hydrogel formulation (10 µg cGAMP hydrogel: P = 0.036 at week 4 and P = 0.047 at week 8; 50 µg cGAMP hydrogel: P = 0.046 at week 4 and P = 0.14 at week 8; **Table S3**). Poly(I:C) incorporation into the hydrogel eventually led to titers 2.8-fold above hydrogel control at week 4 (P = 0.51) and 4.9-fold above hydrogel control at week 8 (P = 0.26), but the titer increase was relatively slow and more heterogeneous than was observed when cGAMP served as the hydrogel adjuvant (**Table S3**). Notably, prior to boosting at week 8, the 10 µg cGAMP hydrogel and 50 µg cGAMP hydrogel titers were over 200-fold (P < 0.0001) and 300-fold (P < 0.0001) greater than their respective bolus controls containing an equivalent cGAMP dose (**Table S4**).

Following administration of a boosting dose at week 8, anti-RBD titers were reliably detected in each bolus delivery group, with 10 μg and 50 μg cGAMP increasing 12-week titers above alum bolus levels by 2.5-fold (P = 0.31) and 7.6-fold (0.0003), respectively (**Table S5**). The advantage of cGAMP adjuvanted alum hydrogels was again apparent following the boost, where incorporation of 10 μg or 50 μg of cGAMP increased post-boost responses by 5-fold (P = 0.031) or 8.5-fold (P = 0.0015), respectively, above the control hydrogel at week 12 (**Table S3**). In contrast, inclusion of poly(I:C) into the hydrogel provided no significant benefit at week 12. While the 50 µg cGAMP hydrogel did not show a significant improvement over the 50 µg poly(I:C) control following the prime dose, the cGAMP group did have a significant advantage after the boost (7.7-fold; P = 0.0046; **Table S4**). Additionally, titers following vaccination with the 50 µg cGAMP hydrogel showed less spread among mice compared to the poly(I:C) hydrogel titers. The average standard deviation across timepoints was 1.4-fold greater and 1.7-fold greater prior to- and following boosting for the poly(I:C) hydrogel compared to the cGAMP hydrogel. Ultimately, cGAMP hydrogel groups achieved anti-RBD titers of 8.8×10^5^ and 1.5×10^6^ for the 10 and 50 µg cGAMP doses, respectively, which represent over 100-fold (P < 0.0001) and 50-fold (P < 0.0001) increases in titer for respective hydrogel groups compared to dose-matched bolus administration (**Table S4**).

We next evaluated the isotypes of IgG that contributed to overall titers at week 12, which serve as an indicator of how each adjuvant was influencing immune signaling. Specifically, elicitation of IgG1 is associated with Th2 dominated immune responses, while IgG2c is associated with Th1-mediated skewing.^[42]^ Alum and the hydrogel system are independently associated with strong, Th2-skewed humoral immunity, and we observed that the major isotype elicited in all vaccine groups was indeed IgG1 (**Figure 2**c).^[31]^ ^[43]^ Inclusion of 10 μg or 50 μg cGAMP in the hydrogel further elevated IgG1 titers by 4.9-fold (P = 0.0006) or 8.5-fold (P < 0.0001), respectively (**Table S6**). While IgG2c titers were generally lower than those of IgG1, IgG2c titers were boosted in a cGAMP dose-dependent manner. As compared to hydrogel control, adding cGAMP at 10 μg or 50 μg increased IgG2c titers by ∼20-fold (P = 0.035) and ∼200-fold (P = 0.0002) (**Figure 2**d; **Table S6**), therefore elevating the relative IgG2c/IgG1 ratio towards a more balanced response (**Figure 2**e). This observation is not surprising as numerous groups have described a role for cGAMP in balancing Th1/Th2 responses, predominantly through (i) increasing inflammatory cytokine signaling and (ii) eliciting cellular immunity which can be a crucial partner of humoral immunity in clearing certain infections.^[26]^ ^[44]^

To confirm that the anti-RBD antibodies elicited from subunit vaccination cross-react with RBD presented in the context of native SARS-CoV-2 spike protein trimers, we then measured anti-spike IgG titers. Additionally, we sought to determine the degree to which antibody titers were influenced by residue substitutions in the spike protein associated with emerging SARS-CoV-2 variants of concern, including B.1.1.7 (Alpha, UK), B.1.351 (Beta, South Africa), and B.1.617.2 (Delta). Across groups, average endpoint titers against the native spike trimer, B.1.1.7 variant and B.1.617.2 variant were largely similar, but decreased to a larger degree for the B.1.351 strain, which is understood to be an immune escape variant.^[45]^ The fold decrease in titer against native trimer compared to the B.1.351 variant was only 1.9 (P = 0.033) for the 10 µg cGAMP hydrogel while it was 4.1 (P < 0.0001) for the bolus equivalent and 11.8 (P = 0.007) for the poly(I:C) hydrogel (**Figure 2**f; **Table S7**). Notably, although titers decreased slightly against both the Alpha and Beta variants for the 10 and 50 µg cGAMP hydrogel groups (along with all other groups), titers remained well above the comparable titers for the native trimer observed for bolus vaccine controls (127- and 59-fold, respectively), demonstrating that hydrogel vaccines provided an improved humoral response compared to bolus controls, even against novel variants of concern (**Figure 2**f).

### 2.3. cGAMP increases neutralizing antibody titers when added as an adjuvant to RBD hydrogel vaccines

We then sought to determine if the elevated antibody titers found in GAMP-adjuvanted hydrogel groups translated to differences in neutralizing activity of the sera. We employed a reporter lentivirus pseudotyped with SARS-CoV-2 spike and measured serum-mediated inhibition of viral entry into HeLa cells overexpressing ACE2.^[46]^ ^[47]^ We first surveyed 12-week sera from all vaccine conditions at a single dilution of 1:250 (**Figure 3**a). Sera from bolus vaccinated groups was found to have no or minimal effect on viral infectivity. More appreciable neutralizing activity was observed in sera from most animals in hydrogel vaccine control and poly(I:C) hydrogel vaccination groups, whereby infectivity decreased to ∼30% of total on average, although heterogeneity in the response was notable. In contrast, significant neutralizing activity was found in sera from all animals vaccinated with cGAMP adjuvanted hydrogels, with average relative infectivity reduced to just 1-3% of total.

**Figure 3.**
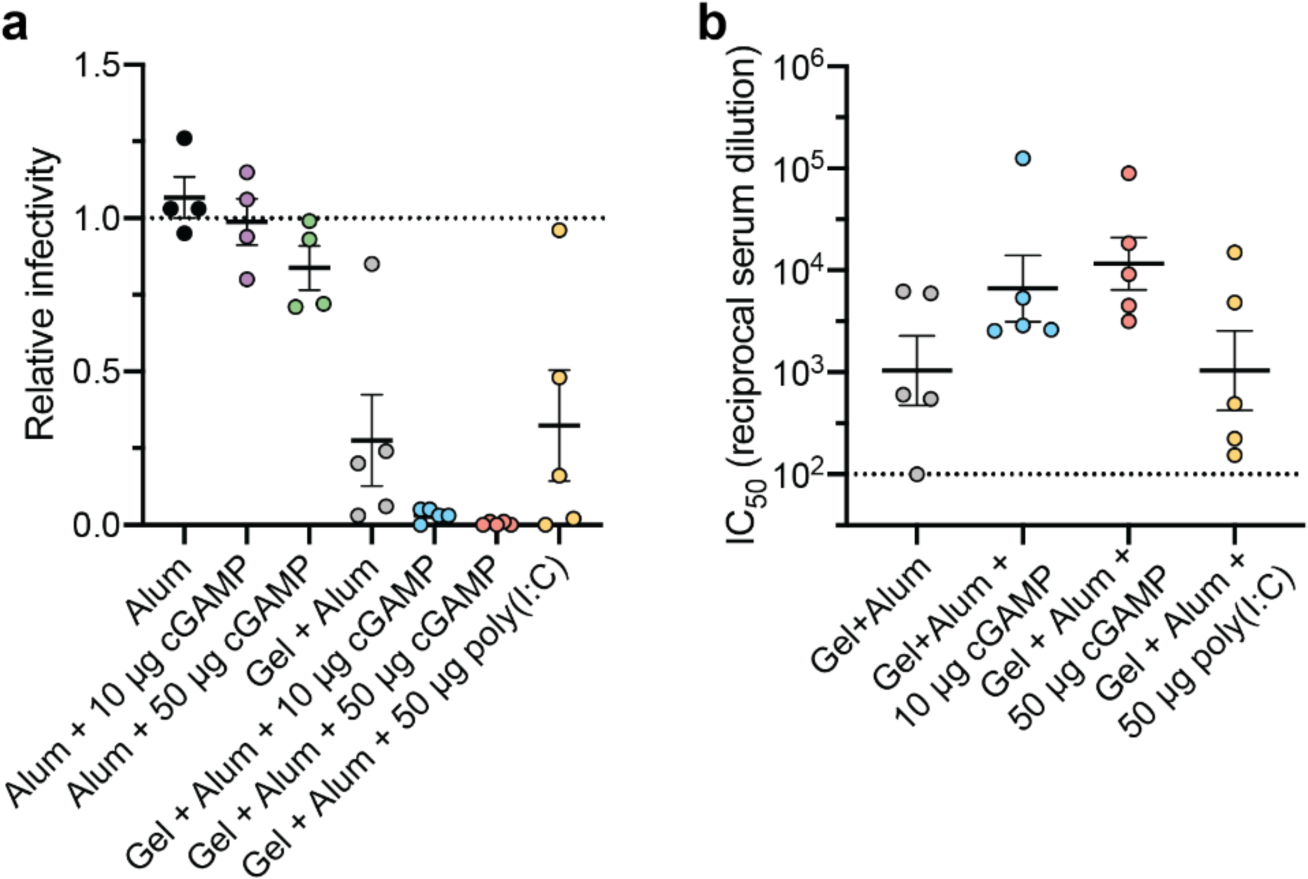
cGAMP-adjuvanted hydrogels provide greater neutralizing antibody titers than bolus and non-adjuvanted controls. (A) Percent infectivity across treatment groups at a 1 to 250 serum dilution. (B) IC50 values determined from neutralization curves **(Figure S2**a-d). Samples with minimal or undetectable neutralizing activity at the 1 to 250 dilution (A) are excluded. (A-B) Data are shown as mean +/- SEM. P values listed in the text were determined using an Ordinary one-way ANOVA with Dunnett’s multiple comparisons test to compare IC50 values (**Table S8**).

We then sought to further quantify the neutralizing potency by assaying a range of sera concentrations from the hydrogel groups in which we observed significant inhibition of infectivity. We determined that the half maximal inhibition of infectivity (IC50) for control- and poly(I:C)-hydrogel groups occurred at reciprocal dilutions of 2.7×10^3^ and 4.1×10^3^, respectively (**Figure 3**b, **Figure S2**a-d). In comparison, sera from hydrogel groups adjuvanted with 10 μg or 50 μg cGAMP had IC50 values of 2.8×10^4^ (P = 0.24 compared to Gel+Alum control) and 2.5×10^4^ (P = 0.098 compared to Gel+Alum control) reciprocal dilutions, respectively, corresponding to an order of magnitude increase in neutralizing activity relative to the control hydrogel group (**Table S8**). Taken together, these data demonstrate that (i) PNP hydrogel-based vaccine administration provides significant benefit for elicitation of humoral immunity versus bolus administration and (ii) adjuvanting hydrogels with cGAMP further boosts elicitation of functional, protective antibodies. We also compared our neutralization results to other COVID-19 mouse immunization studies by plotting a set of IC50 values found in the literature (**Figure S3**, **Table S9**).^[31]^ ^[48]^ ^[49]^ ^[50]^ ^[51]^ ^[52]^ ^[53]^ Our 50 µg cGAMP hydrogel containing wildtype RBD resulted in IC50 values well over 10^4^, greatly exceeding the FDA “high titer” cutoff of IC50∼10^2.4^.^[54]^ Notably, the 50 µg cGAMP hydrogel IC50 value is comparable to other leading subunit vaccine candidates containing RBD or Spike constructs found in the literature (**Figure S3**, **Table S9**).^[31]^ ^[48]^ ^[49]^ ^[50]^ These results provide proof of principle that controlled release of cGAMP from hydrogel-based vaccines represents an effective strategy to unleash the adjuvant activity of a molecule with a notoriously challenging pharmacokinetic profile.

### 2.4. cGAMP promotes migration of numerous immune cell types to the hydrogel vaccine niche

We previously found that PNP hydrogels created a physical inflammatory niche into which cells can infiltrate.^[32]^ Moreover, the antigen and adjuvant cargo crucially shape the nature of the inflammatory microenvironment formed within the gel. We therefore evaluated how addition of cGAMP influences the degree and composition of cellular infiltrate into the hydrogel. To evaluate the specific cGAMP-related cell infiltration and activation, we used poly(I:C) as an alternative type I IFN inducer. Four days following *in vivo* subcutaneous injection, hydrogels were excised and then the total cellular infiltration was quantified via flow cytometry (**Figure 4**a / **S4**). The addition of cGAMP increased the average infiltrating cell number in the gel (**Figure 4**b), with some heterogeneity in the magnitude of the effect observed across animals. Similarly, the amount of infiltrating CD45^+^ leukocytes also increased upon addition of cGAMP (**Figure 4**b). The same holds true for poly(I:C), where live cells and especially leukocytes increased as well (**Figures S5**).

**Figure 4.**
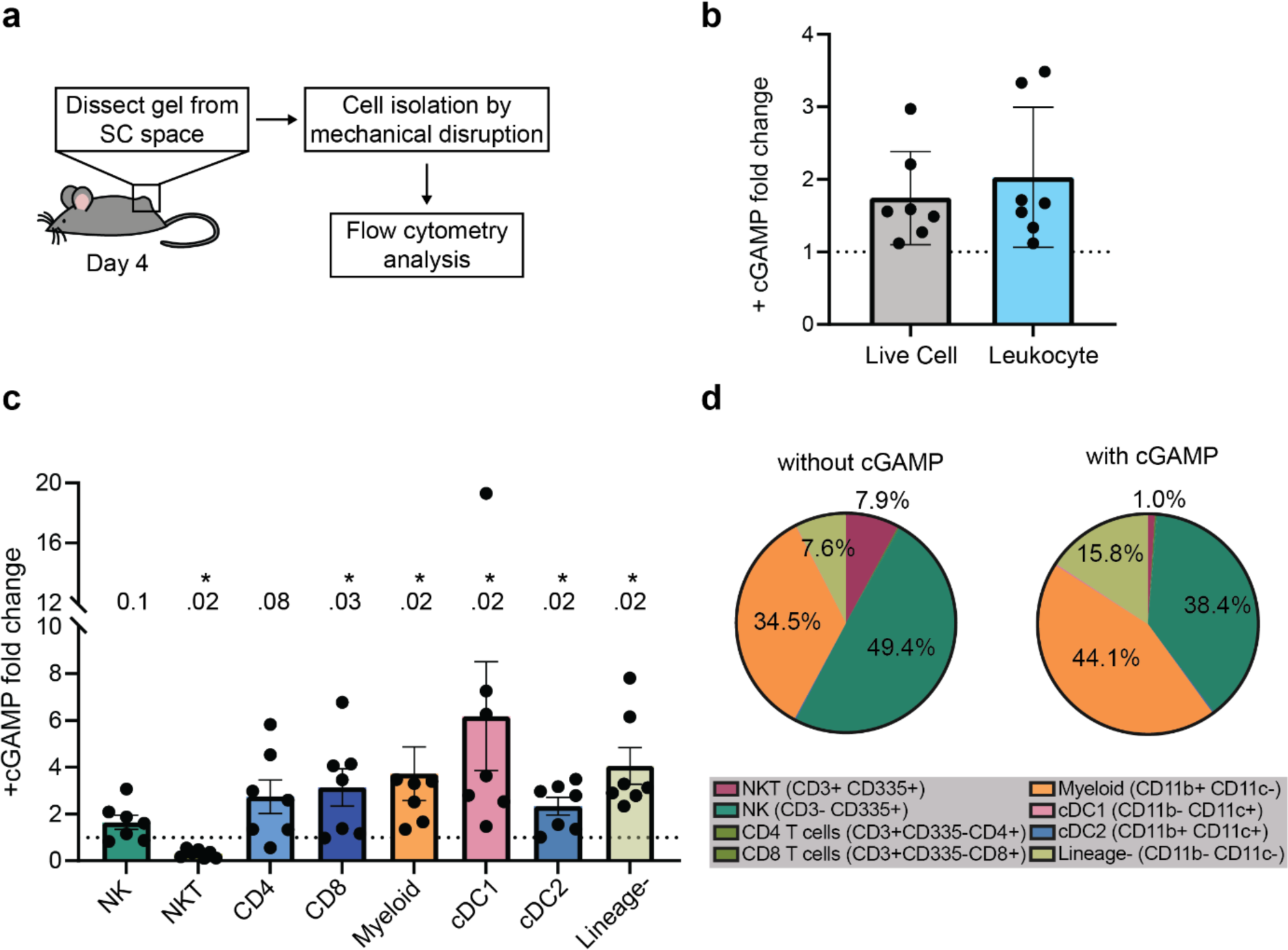
Characterization of cellular infiltrate to the hydrogel vaccine niche. (A) Schematic of the experimental workflow: four days following injection of hydrogels containing RBD, alum, and the absence or presence of 10 µg cGAMP, gels were excised and dissociated. Total cellular infiltrate was then analyzed by flow cytometer. (B) Fold change of flow cytometry counts of live cells and CD45^+^ leukocytes compared to hydrogels without cGAMP. (C) Fold change of proportions of major immune subsets among total CD45^+^ infiltrate compared to hydrogels without cGAMP. (D) Percentages of major immune subsets among total CD45+ infiltrate, with or without 10 µg cGAMP. For (B-C), n = 6 (-cGAMP) and n = 7 mice (+ cGAMP) with error bars representing mean ± SEM; p values are determined by a Wilcoxon signed-rank test.

Addition of cGAMP to hydrogels resulted in a shift in several immune subsets compared to control hydrogels containing alum and RBD alone (**Figure 4c**). Among lymphoid cell populations infiltrating the gel, cytotoxic T lymphocytes CD8^+^ cells were elevated and NKT cells were downregulated in their recruitment to hydrogels with cGAMP compared to control hydrogels. The infiltration of NK cells and T helper CD4^+^ cells had a more subtle response and were non-significantly yet slightly elevated. Additional cell subsets which showed significant changes with cGAMP inclusion within the gels were myeloid cells and dendritic cells, both conventional type 1 and type 2 dendritic cells (cDC1 and cDC2, respectively). cDC1s cross-present exogenous antigen on MHC I for activation of cytotoxic T lymphocytes, which are instructed to kill infected cells. cDC2s, while also able to cross-present, present their antigen on MHC II molecules and are thus the main activators of T helper cells which in turn are crucial for activation of both CTL and the humoral response mediated by antibody-producing B cells.^[13]^ ^[21]^ In examining how the percent representation of each subset shifted among the total number of infiltrating cells (**Figure 4d**), the most pronounced effect of cGAMP appeared to be a skewing towards myeloid cells and the prevalence of a CD45+ but CD19-CD3-CD335-CD11b-CD11c-lineage negative subset, the identification of which will require further surface markers to the flow cytometry panel in future studies. Overall, these results suggest that cGAMP plays an important role in affecting cellular migration to the site of the hydrogel depot, which we predict contributes to downstream benefits in antigen presentation and cellular activation that enable the enhanced humoral responses we observed in these vaccine groups.

## 3. Discussion

The efficacy of subunit vaccines is determined by the selection of antigen, adjuvant(s), and the spatiotemporal control of their exposure to the immune system, all of which crucially interact to shape the resulting immune response. Both the RBD as an antigen and cGAMP as an adjuvant offer promising advantages, but each also possesses significant limitations related to ineffective delivery and poor localization within immune compartments. Here, we demonstrated that an injectable, self-assembled PNP hydrogel system simultaneously helps overcome delivery challenges posed by RBD and cGAMP, and that co-administration of these agents in this easily produced vaccine format leads to significant and functionally important increases in immunogenicity. The approach used in our study is well suited for the development of WT or variant-specific COVID-19 vaccines. The RBD protein, as opposed to the full spike trimer, is easily produced in large quantities, and is more stable to physical and chemical degradation compared to the spike protein and mRNA vaccines. Moreover, while alum is a widely used adjuvant for vaccines, the hydrogel components reported here constitute new chemical entities and will have to be shown to be safe in clinical trials as part of a complete vaccine drug product review. Furthermore, cGAMP is an unmodified biomolecule with ENPP1 as its abundant degradation enzyme, which will only exert its function in and immediately around the gel niche before being degraded, avoiding systemic inflammation. A potential limitation could be a robust cellular response, which we did not investigate here. With the onset of new serotypes, SARS-CoV-2 is here to stay and there will be a continuous need for adaptable, cheap, stable, and easy to produce vaccines for the world population in the coming years.

While the importance of targeting the RBD is illustrated by the high fraction of neutralizing antibodies that target the RBD, we and others have now shown that engineering of delivery systems for the protein domain or its multimerization are required to generate appreciable anti-RBD antibody responses in vaccination.^[9]^ ^[31]^ ^[51]^ For example, we have shown previously that delivery of subunit vaccines from a PNP hydrogel depot improves the humoral response compared to a dose-matched bolus control likely because prolonged antigen release better mimics the kinetics of antigen exposure occurring during a natural infection.^[31]^ ^[32]^ It is noteworthy that the levels of neutralizing IgG antibodies against the Delta variant spike protein are on the same level as for the WT spike, the RBD of which we used in our hydrogel. This is contrasting findings in both mouse and human, where the titers against Delta compared to WT were significantly lower^[55]^ ^[56]^. A brief literature search showed that IC50 values from neutralization studies following immunization with our cGAMP hydrogel vaccine greatly exceeded the FDA “high titer” mark and exceeded IC50 values for both Spike- and RBD-encoding mRNA vaccines in mice. It is important to note that there were differences in dosing, timelines, and neutralization assay procedures across these studies and that IC50 values are an indication of the humoral response and not the cell-mediated response, which is known to also play a role in protection from SARS-CoV-2.

A main finding from this work is that our PNP hydrogel is a suitable delivery system for cGAMP as a subunit vaccine adjuvant. Within the context of a COVID-19 vaccine, cGAMP is particularly useful as an adjuvant because of its role in inducing potent cell-mediated (Th1) responses. Previous studies have shown that less severe cases of SARS-CoV-1 were associated with rapid onset of a Th1 response, while Th2 responses were linked to greater lung inflammation and worse outcomes.^[57]^ Although the PNP hydrogel itself was previously found to skew towards a humoral (Th2) response, we observe here that addition of cGAMP inverted this effect in a dose-dependent manner.^[31]^ ^[32]^ Notably, the average IgG2c/IgG1 ratio for the 50 µg cGAMP hydrogel group was higher than that of the dose-matched bolus group, suggesting that hydrogel was potentiating cGAMP’s ability to skew towards Th1. Future studies will reveal the degree to which cGAMP-embedded gels affect Th1 skewing at the level of cellular function, including induction of antigen-specific cytotoxic CD8^+^ T cells.

Another striking finding was that cGAMP clearly outperformed poly(I:C) as an adjuvant within the hydrogel depot, with poly(I:C) providing no additional benefit after the boosting vaccine dose versus control gel. We speculate that several factors enable cGAMP to more rapidly, consistently, and robustly adjuvant the hydrogel-based vaccine versus poly(I:C). As a small molecule, cGAMP is more likely to be evenly distributed throughout the assembled matrix and diffuse more quickly within and out of the gel. While extracellular cGAMP must enter cells to activate STING in the cytosol, we and others recently established that cells possess specific importers that allow for the small molecule’s uptake.^[27]^ ^[28]^ ^[29]^ ^[30]^ In contrast, poly(I:C) is very large (1.5 – 8 kb) and is imported through endosomal uptake. Poly(I:C) then can activate toll-like receptor 3 within endosomes or can escape to the cytosol to activate RIG-I like receptors.^[58]^ The different routes of uptake and pattern recognition receptor activation may also explain the differences we observed between cGAMP and poly(I:C) in our cell analysis. The PNP hydrogel system was designed so that diverse antigen cargo can be easily loaded into it, and our results suggest that incorporation of cGAMP into subunit vaccine hydrogels of many different antigens could represent a flexible strategy for production of vaccines against other viral and infectious agents, especially those that require more durable immunity with balanced Th1/Th2 responses for clearance.

We have previously shown that the hydrogel delivery system provides sustained release of both antigens and adjuvants *in vivo*.^[31]^ ^[32]^ Ideally, we would have liked to perform thorough cGAMP biodistribution analyses here to verify that the cGAMP hydrogel groups achieved high titers and neutralization through this same mechanism; however, two properties of cGAMP make these studies challenging. Since cGAMP is a small cyclic dinucleotide, it is difficult to label for tracking studies while maintaining native-like properties and activity. Moreover, cGAMP is rapidly degraded in circulation by ENPP1, which precludes direct detection of systemic cGAMP levels unless ENPP1 inhibitors are continually dosed at elevated concentrations.^[14]^ ^[15]^ While we do not present *in vivo* release or biodistribution data for cGAMP, in previous work we showed that RBD was released from the hydrogel *in vivo* with a half-life of ∼5.5 days, which is within a similar window to the half-life of ∼3.5 days for cGAMP release observed here in our *in vitro* study.^[31]^

Our initial examination of the hydrogel infiltrate revealed that cGAMP is playing a major role in promoting immune cell recruitment to the inflammatory niche at the site of vaccination. The site of administration is known to influence the quantity and phenotypes of tissue-resident immune cells that initially interact with the vaccine, and thereby strongly affects the magnitude, duration and quality of the adaptive immune responses.^[5]^ ^[59]^ Yet, while this relationship is known to exist, the exact mechanisms have not yet been well studied in the context of vaccination.^[5]^ ^[59]^ The effect of cGAMP on increasing immune infiltration to hydrogels containing alum and RBD was significantly more profound than the role played by alum and RBD compared to empty hydrogels. These results suggest that cGAMP plays an early and crucial role in immune activation, as would be desired of an effective adjuvant. The elevated recruitment of various cell types, including T-helper cells, cytotoxic T cells, cDC1 and cDC2 cells, give us initial insight as to how cGAMP as an adjuvant may work on a cellular and systemic level. It is worth noting that these data at present do not reveal whether cGAMP is preferentially interacting with cells localized around the hydrogel, or if cGAMP that reaches the draining lymph node serves to mobilize cells to traffic to the hydrogel. It is however, likely that cGAMP only acts over short distances on account of rapid degradation by ENPP1 in the tissue.^[14]^ ^[16]^ Future detailed biological studies are expected to shed light on the central mechanisms by which hydrogel-controlled cGAMP delivery affects innate immune cell activation state, presentation of co-delivered antigen, and the development of adaptive cellular effectors for both humoral and cell-mediated protection. While the hydrogel-based vaccines elicited antisera with appreciable *in vitro* neutralizing activity, assessing the degree of induced protection *in vivo* through viral challenge studies remains an important aspect for future work to develop cGAMP-adjuvanted subunit hydrogel vaccines for SARS-CoV-2 and beyond.

## 4. Conclusions

Overall, our data demonstrate that delivery of cGAMP from a PNP hydrogel containing the commonly used adjuvant, alum, and the critical SARS-CoV-2 antigen, RBD, results in significantly higher anti-RBD antibody titers and improved neutralization as compared to both dose-matched bolus controls and the hydrogel vaccine lacking cGAMP. Immunization with the cGAMP hydrogel vaccines containing RBD resulted in higher cross-reactive antibody titers against the recent SARS-CoV-2 spike protein variants Alpha, Beta, and Delta. As an early indication of cGAMP’s ability to act as an effective adjuvant in this system, we found that inclusion of cGAMP in the hydrogel increases infiltration of key immune populations into the hydrogel vaccine niche as early as 4 days following injection. These results support the use of PNP hydrogels as a means to overcome the delivery limitations of cGAMP and illustrate that cGAMP, when delivered in this manner, is an effective adjuvant for improving immunogenicity against SARS-CoV-2 RBD antigen. Given the findings presented here, cGAMP-adjuvanted hydrogels represent an effective and scalable subunit vaccine platform that could be readily adapted and evaluated for use against a broad range of infectious agents.

## 4. Experimental Section

### 4.1. Materials

#### Mammalian Cell culture

Expi293F cells (ThermoFisher) were maintained in a 2:1 mix of FreeStyle293:Expi293 Expression medium (ThermoFisher) and grown at 37 °C and 8% CO2 while shaking at 120 rpm. HEK293T cells (ATCC) were maintained in DMEM (Cellgro) supplemented with 10% FBS, L-glutamate, 1% penicillin-streptomycin, and 10 mM HEPES. ACE2/HeLa cells were generously provided by Dennis Burton and were maintained in DMEM (Cellgro) supplemented with 10% FBS, L-glutamate, 1% penicillin-streptomycin, and 10 mM HEPES.^[47]^

#### Chemicals

HPMC (meets USP testing specifications), N,N-diisopropylethylamine (Hunig’s base), hexanes, diethyl ether, N-methyl-2-pyrrolidone (NMP), dichloromethane (DCM), lactide (LA), 1-dodecylisocynate, and diazobicylcoundecene (DBU) were purchased from Sigma-Aldrich and used as received. Monomethoxy-PEG (5 kDa) was purchased from Sigma-Aldrich and was purified by azeotropic distillation with toluene prior to use. For in vitro studies, 2’3’-cyclic-GMP-AMP (cGAMP) was synthesized and purified in-house as previously described.^[27]^ For *in vivo* studies, vaccine-grade 2’3’-cyclic-GMP-AMP (cGAMP) was purchased from Invivogen.

#### Plasmids

The SARS-CoV-2 RBD DNA construct was kindly provided by Dr. Florian Krammer.^[60]^ The expression plasmid (pCAGGS) contains a CMV promoter, followed by the native SARS-CoV-2 Spike signal peptide (residues 1-14), RBD-encoding residues 319-541 from the SARS-CoV-2 Wuhan-Hu-1 genome sequence (GenBank MN908947.3), and a C-terminal hexa-histidine tag. A five-plasmid system was used for production of SARS-CoV-2 spike pseudotyped lentivirus: the lentiviral packaging vector (pHAGE_Luc2_IRES_ZsGreen), the SARS-CoV-2 spike (sequence from Wuhan-Hu-1 strain of SARS-CoV-2, GenBank NC_045512), and lentiviral helper plasmids (HDM-Hgpm2, HDM-Tat1b, and pRC-CMV_Rev1b).^[46]^

### 4.2. Methods

#### Expression and purification of SARS-CoV-2 Spike RBD

The mammalian expression plasmid for RBD production was a kind gift from Dr. Florian Krammer and was previously described in detail.^[48]^ Transient transfection of Expi293F cells was performed at a density of ∼3-4 x 10^6^ cells mL^-1^. Per liter of transfected cells, 568 µg DNA in 113 mL culture medium was complexed with 1.48 mL FectoPro (Polyplus) for 10 min at RT. The transfection mixture was then added to cells along with boosting D-glucose (4 g L^-1^ final) and 2-propylpentanoic (valproic) acid (3 mM final). Culture supernatants were harvested 3-5 days post-transfection via centrifugation at 7,000 x g for 15 minutes followed by 0.22 µm filtration. Samples were snap frozen and stored at -20 °C until use.

HisPur NiNTA resin (ThermoFisher) was washed 3x with ∼10 column volumes wash buffer (10 mM imidazole in 1X PBS, pH 7.4). Culture supernatant was diluted 1:1 (10 mM imidazole in 1x PBS, pH 7.4) and incubated with resin at 4 °C with gentle stirring. Resin/supernatant mixture was then loaded into gravity flow columns and washed 1x with wash buffer. Protein was eluted with 250 mM imidazole in 1x PBS and spin concentrated (10 kDa MWCO, Amicon). Size-exclusion chromatography was performed using a GE Superdex 200 Increase 10/300 GL column pre-equilibrated in 1x Dulbecco’s phosphate-buffered saline (Gibco). Fractions were evaluated by SDS-PAGE (4-20% Mini-PROTEAN TGX, ThermoFisher), pooled, and spin concentrated. Purified RBD was supplemented with 10% glycerol, filtered through a 0.22 µm filter, snap frozen, and stored at -20 °C.

#### Preparation of HPMC−C12

HPMC−C12 was prepared according to previously reported procedures.^[32]^ ^[61]^ HPMC (1.0 g) was dissolved in NMP (40 mL) by stirring at 80 °C for 1 h. Once the solution cooled to RT, 1-dodecylisocynate (105 mg, 0.5 mmol) and N,N-diisopropylethylamine (catalyst, ∼3 drops) were dissolved in NMP (5.0 mL). This solution was added dropwise to the reaction mixture, which was then stirred at RT for 16 h. This solution was then precipitated from acetone, decanted, re-dissolved in water (∼2 wt %), and placed in a dialysis tube for dialysis for 3−4 days. The polymer was lyophilized and reconstituted to a 60 mg mL^-1^ solution with sterile PBS.

#### Preparation of PEG−PLA NPs

PEG−PLA was prepared as previously reported.^[32]^ ^[61]^ Monomethoxy-PEG (5 kDa; 0.25 g, 4.1 mmol) and DBU (15 μL, 0.1 mmol; 1.4 mol % relative to LA) were dissolved in anhydrous dichloromethane (1.0 mL). LA (1.0 g, 6.9 mmol) was dissolved in anhydrous DCM (3.0 mL) with mild heating. The LA solution was added rapidly to the PEG/DBU solution and was allowed to stir for 10 min. The reaction mixture was quenched and precipitated by 1:1 hexane and ethyl ether solution. The synthesized PEG−PLA was collected and dried under vacuum. Gel permeation chromatography (GPC) was used to verify that the molecular weight and dispersity of polymers meet our quality control (QC) parameters. NPs were prepared as previously reported. ^[32]^ ^[61]^ A 1 mL solution of PEG−PLA in DMSO (50 mg mL^-1^) was added dropwise to 10 mL of water at RT under a high stir rate (600 rpm). NPs were purified by centrifugation over a filter (molecular weight cutoff of 10 kDa; Millipore Amicon Ultra-15) followed by resuspension in PBS to a final concentration of 200 mg mL^-1^. NPs were characterized by dynamic light scattering (DLS) to ensure they passed our QC parameters.

#### PNP Hydrogel Preparation

PNP hydrogels using 2:10 of HPMC−C12: PEG−PLA NP were made by mixing a 2:3:1 weight ratio of 6 wt % HPMC−C12 polymer solution, 20 wt % NP solution, and PBS (with or without adjuvants). The NP and aqueous components were loaded into one syringe, the HPMC-C12 was loaded into a second syringe and components were mixed using an elbow connector. After mixing, the elbow was replaced with a 21-gauge needle for injection.

#### Vaccine Preparation

All the following vaccine variants contained 10 μg SARS-CoV-2 RBD per dose. Each bolus vaccine dose (100 μL injection volume) contained 115 μg Alhydrogel (Invivogen) in PBS with 10 μg or 50 μg 2’3’-cGAMP (Invivogen) added where indicated. PNP hydrogel vaccines were made by diluting respective vaccine cargo in PBS and then mixing a 2:3:1 weight ratio of 6 wt % HPMC−C12 polymer solution, 20 wt % NP solution, and PBS (-/+ cargoes). As indicated, each PNP hydrogel dose (150 μL injection volume) contained 115 μg Alhydrogel, 10 μg or 50 μg 2’3’-cGAMP, or 50 μg Poly(I:C) (Sigma-Aldrich).

#### cGAMP In vitro Release Studies

Hydrogels were prepared as described above in the “PNP Hydrogel Preparation” and “Vaccine Preparation” sections. Glass capillary tubes were plugged at one end with epoxy and 150 µL of gel was injected into the bottom of 3 separate tubes per treatment. 350 µL PBS was added on top of each gel. Tubes were stored upright in an incubator at 37 °C for about 3 weeks. At each timepoint, ∼300 µL of PBS was removed and the same amount was replaced. The amount of cGAMP released at each timepoint was determined by measurement of A260 absorbance relative to a standard curve by Nanodrop. The cumulative release was calculated and normalized to the total amount released over the duration of the experiment. Points were fit with the nonlinear “One site – Total fit” model in GraphPad Prism and the half-life was determined.

#### Mouse immunizations

C57BL/6 (B6) mice were purchased from Charles River and housed at Stanford University. Female mice between 8 and 10 weeks of age at the start of the experiment were used. Mice received a subcutaneous (SC) injection of bolus (150 μL) or hydrogel (150 μL) vaccine on their backs under brief isoflurane anesthesia. PBS injections used a 26-gauge needle, and gel injections used a 21-gauge needle. Blood was collected by tail vein bleeds for survival studies or through cardiac puncture for terminal studies, with serum processed using Z gel microtubes (Sarstedt). In the immune infiltration experiment, SC gels were collected for analysis after euthanasia.

#### Anti-RBD ELISA with mouse serum

MaxiSorp round-bottom immuno 96-well plates (ThermoFisher) were coated with RBD antigen (2 µg mL^-1^ in PBS, pH 7.4, 50 µL well^-1^) overnight at 4 °C. Additional ELISAs were done with spike and spike mutant-coated plates (wildtype, Alpha (UK variant, B.1.1.7), Beta (South Africa variant, B.1.351), Delta (B.1.617.2); Sino Biological). After coating, plates were washed 3x (PBS with 0.05% Tween-20). Plates were blocked with PBS containing 1% BSA (250 µL well^-1^) for 1 h at RT and then washed 3x. Mouse sera were diluted in diluent buffer (PBS with 1% BSA and 0.05% Tween-20) starting at 1:50 and 4-fold serially dilution. A negative reference control consisting of pooled non-immunized sera was included in duplicate on each plate at 1:50 dilution for all determinations. Diluted sera were added to coated/blocked plates (50 µL well^-1^) and incubated for 2 h at RT (for total IgG analysis) or overnight at 4 °C (for IgG1 and IgG2 analysis). Plates were washed 5x. The respective HRP-conjugated secondaries were diluted in dilution buffer and incubated as follows:

- Total IgG: goat anti-mouse IgG Fc, Invitrogen #A16084, 1:10,000, 1 h at RT
- IgG1: goat anti-mouse IgG1 Fc, Abcam #ab97240, 1:50,000, 2 h at 4 °C
- IgG2c: goat anti-mouse IgG2c Heavy chain, Abcam #ab97255, 1:10,000, 1 h at RT).

ELISA plates were then washed 6x. Plates were developed using high-sensitivity TMB ELISA subtrate (Abcam) for 8 min at RT and quenched with 1 M HCl. Absorbance at 450 nm was measured using a Tecan Spark plate reader. The endpoint threshold was set as 2 times the reference negative control average obtained each day. Sample dilution curves were imported into GraphPad Prism 8.4.1, fit with a three-parameter non-linear regression (baseline constrained to 0), and dilution titer value at which the endpoint threshold was crossed for each curve was imputed. Samples failing to meet endpoint threshold at a 1:50 dilution were set to a titer cutoff of 1:25 or below the limit quantitation for the assay.

#### Mouse cytokine ELISAs

Determination of cytokines in mouse sera were determined using the following ELISA kits: IFNα concentrations were determined using PBL Assay Science VeriKine-HS Mouse IFN-α All Subtype ELISA Kit, CXCL10 concentrations were determined using R&D Systems CXCL10/IP-10/CRG-2 DuoSet ELISA Kit, and TNFα concentrations were determined using R&D Systems Mouse TNF-α Quantikine ELISA Kit. The manufacturer recommended protocols were followed with sera assayed at a 1:20 dilution and then concentrations imputed from standard curves.

#### SARS-CoV-2 pseudotyped lentivirus production and viral neutralization assays

SARS-CoV-2 spike pseudotyped lentivirus was produced in HEK293T cells via calcium phosphate transfection. Six million cells were seeded in D10 medium (DMEM + additives: 10% fetal bovine serum, L-glutamate, penicillin, streptomycin, and 10 mM HEPES) in 10-cm plates one day prior to transfection. A five-plasmid system was used for viral production consisting of the lentiviral packaging vector (pHAGE_Luc2_IRES_ZsGreen), the SARS-CoV-2 spike, and lentiviral helper plasmids (HDM-Hgpm2, HDM-Tat1b, and pRC-CMV_Rev1b).^[46]^ The spike vector contained the full-length wild-type spike sequence from the Wuhan-Hu-1 strain of SARS-CoV-2 (GenBank NC_045512). Plasmids were added to filter-sterilized water as follows: 10 µg pHAGE_Luc2_IRS_ZsGreen, 3.4 µg SARS-CoV-2 spike, 2.2 µg HDM-Hgpm2, 2.2 µg HDM-Tat1b, and 2.2 µg pRC-CMV_Rev1b in a final volume of 500 µL. HEPES-buffered Saline (2X, pH 7.0) was added dropwise to this mixture to a final volume of 1 mL. To form transfection complexes, 100 µL 2.5 M CaCl2 were added dropwise while the solution was gently agitated. Transfection reactions were incubated for 20 min at RT, then added dropwise to plated cells. Medium was removed ∼24 h post transfection and replaced with fresh D10 medium. Virus-containing culture supernatants were harvested ∼72 h post transfection via centrifugation at 300 x g for 5 min and filtered through a 0.45 µm filter. Viral stocks were aliquoted and stored at -80°C until use. For viral neutralization assays, ACE2/HeLa cells were plated in white-walled clear-bottom 96-well plates at 5,000 cells well^-1^ 1 day prior to infection.^[47]^ Mouse serum was centrifuged at 2,000 x g for 15 min, heat inactivated for 30 min at 56 °C and diluted in D10 medium. Titered virus was diluted in D10 medium, added to diluted sera, and incubated for 1 h at 37°C. Virus:sera dilutions were then transferred to the plated ACE2/HeLa from which seeding media had been aspirated. Polybrene was then spiked in to each well for a final concentration of 5 µg mL^-1^ and plates were incubated at 37 °C for ∼48 h. Cells were lysed by adding BriteLite assay readout solution (Perkin Elmer) and luminescence values were measured with a Tecan Spark plate reader. Each plate was normalized by averaging RLUs from wells with cells only (0% infectivity) and virus only (100% infectivity). Normalized values were fit with a three-parameter non-linear regression inhibitor curve in GraphPad Prism 8.4.1 to obtain IC50 values. Fits for all serum neutralization assays were constrained to have a value of 0% at the bottom of the fit. The limit of quantitation for this assay is approximately 1:100 serum dilution. Serum samples that failed to neutralize or that neutralized at levels higher than 1:100 were set at the limit of quantitation.

#### Flow cytometry analysis of immune cell infiltration

Gels were injected as described in *mouse immunizations*. After four days, mice were euthanized, and gels, draining inguinal lymph nodes and spleens were excised and placed in 10 mL of media. Subsequently, gels were placed in a 5 cm Petri dish and mechanically disrupted between 2 frosted glass slides. The resulting suspension was strained through a 100 μM filter into a 50 mL conical tube and washed 2x with PBS. Lymph nodes and spleens were incubated in RPMI supplemented with 10% FBS containing 20 μg / ml DNase I type IV (Sigma-Aldrich) and 1 mg / ml collagenase from Clostridium histolyticum (Sigma-Aldrich) at 37 °C for 20 min. Organs were passed through a 100-μm cell strainer (Sigma-Aldrich), and red blood cells were lysed with red blood cell lysis buffer (155 mM NH4Cl, 12 mM NaHCO3 and 0.1 mM EDTA) for 5min at room temperature. Cell pellets (gel, lymph node, and spleen ∼200 μL in PBS) were then transferred to a 96-well plate for staining. Samples were first stained with LIVE/DEAD Fixable Blue Dead Cell Stain (Invitrogen) for 30 min then fixed and permeabilized with eBioscience Foxp3/Transcription Factor Staining Buffer Set (Invitrogen). Samples were next Fc-blocked for 10 min using TruStain fcX (BioLegend), and then stained for 1 h (see **Supplemental Table S10** for antibodies and dilutions). Flow cytometry analysis for this project was done on an Aurora analyzer (Cytek) in the Stanford Shared FACS Facility.

#### Animal Protocol

All animal studies were performed in accordance with National Institutes of Health guidelines and with the approval of the Stanford Administrative Panel on Laboratory Animal Care (Protocol APLAC-32109).

#### Statistical Analysis

Statistical analyses were conducted using GraphPad Prism. Comparisons of titer data between treatment groups and comparisons of titers to different spike variants within a treatment group were done using 2way ANOVAs with either Dunnett’s or Tukey’s multiple comparisons test (depending on number of groups compared) on the logged titer values (**Figure 2**, **Table S3-7**). Comparisons of IC50 values were done using an Ordinary one-way ANOVA with Dunnett’s multiple comparisons test (**Figure 3**, **Table S8**). Comparisons of fold-change data were done using a one sample Wilcoxon signed-rank test comparing to a hypothetical fold change of 1 (**Figure 4**). Statistical analysis comparing cytokine levels were done using an ordinary one-way ANOVA with Tukey’s multiple comparisons test (**Tables S1** and **S2**). Select P values are shown in the text and remaining P values are in **Tables S1-8**.

## Materials and Data Availability

All data needed to evaluate the conclusions in the paper are present in the paper and/or the Supplementary Materials. Further information and requests for resources or raw data should be directed to and will be fulfilled by the lead contact, Eric Appel.

## Acknowledgements

We would like to thank all members of the Appel and Li labs for their useful discussion and advice throughout this project. We want to acknowledge the staff of the BioE/ChemE Animal Facility who cared for our mice. Flow cytometry analysis for this project was done on an Aurora analyzer (Cytek) in the Stanford Shared FACS Facility. This research was financially supported by the Center for Human Systems Immunology with the Bill and Melinda Gates Foundation (OPP1113682; OPP1211043). This work was also supported by the Stanford Maternal and Child Health Research Institute postdoctoral fellowship (to AEP), the Stanford School of Medicine Startup fund (to LL), and Stanford Medical School Discovery Innovation Fund (to LL). We thank Dr. Jesse Bloom, Kate Crawford, Dr. Dennis Burton, and Dr. Deli Huang for sharing the plasmids, cells, and invaluable advice for implementation of the spike-pseudotyped lentiviral neutralization assay. We thank the NIH Cell and Molecular Biology Training Program (T32 GM007276; to ECG) and the Eastman Kodak Fellowship (BSU).

