## Supplementary Information for "A cGAMP-containing hydrogel for prolonged SARS-CoV-2 RBD subunit vaccine exposure induces a broad and potent humoral response"

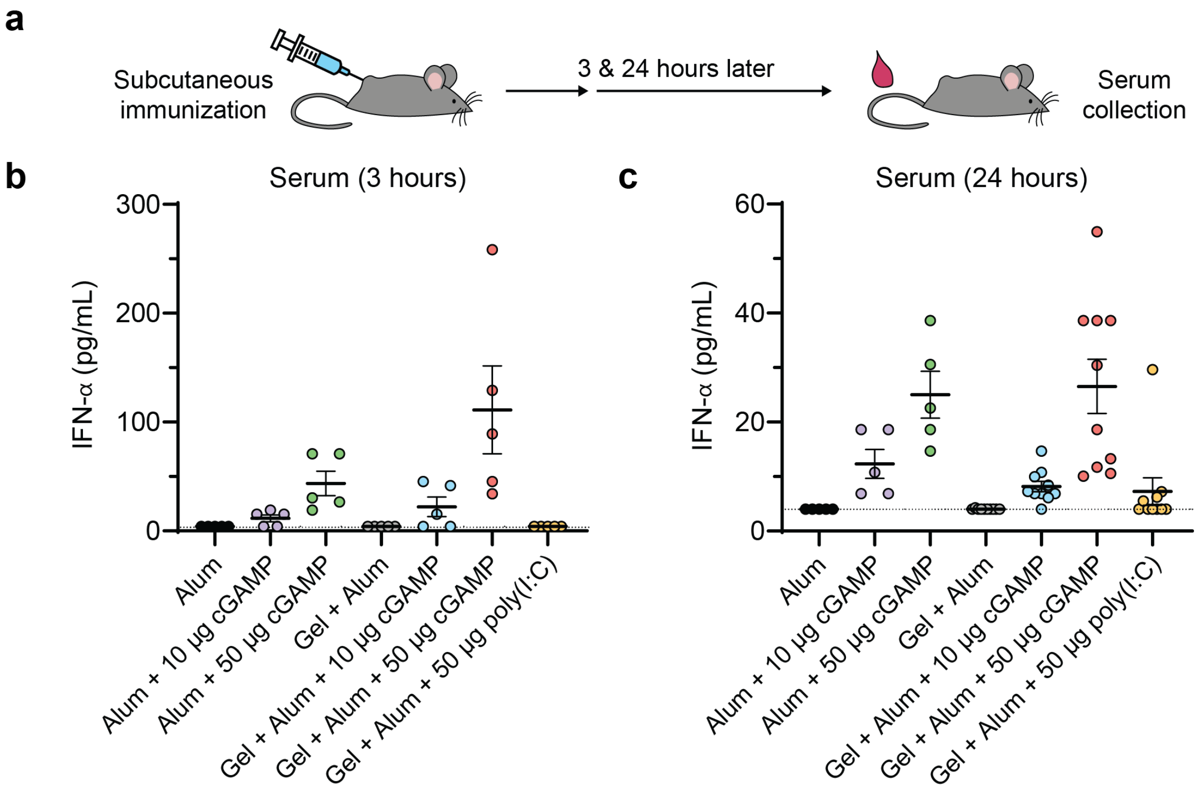


**Figure S1**. (A) Schematic showing mice were immunized by subcutaneous injection and serum was collected for IFNα quantification 3 and 24 hours later. (B-C) IFNα concentrations in serum as determined by ELISA at 3 hours (B) and 24 hours (C) after immunization. Data are shown as mean +/- SEM.

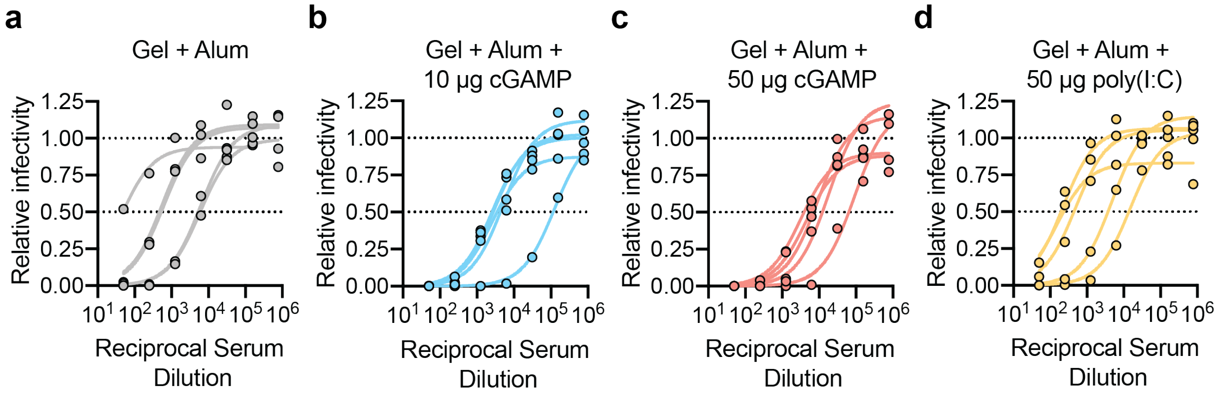


**Figure S2**. (A-D) Relative infectivity across a range of serum dilutions as determined by a SARS-CoV-2 spike-pseudotyped viral neutralization assay for all treatment groups that showed a decrease in infectivity at the 1:250 serum dilution: Gel + Alum (A), Gel + Alum + 10 µg cGAMP (B), Gel + Alum + 50 µg cGAMP (C), and Gel + Alum + 50 µg poly(I:C) (D).

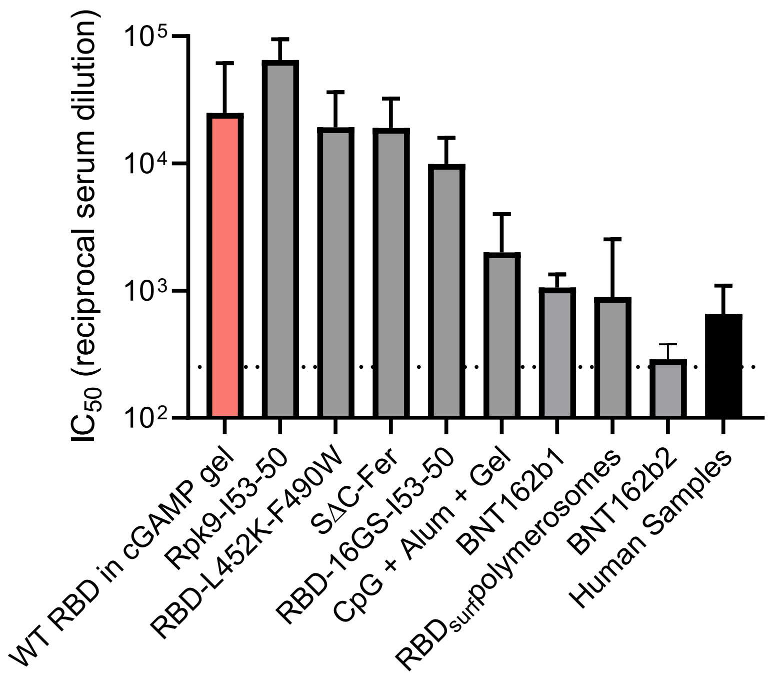


**Figure S3**. Comparison of $IC_{50}$ values following immunization of mice with various RBD or spike constructs. Human convalescent serum data includes samples collected from patients 4-10 weeks after symptom resolution. Human data are compiled from two publications.[1] [2] Data are shown as mean + s.d. Data were extracted from published plots using the following data extraction software: https://apps.automeris.io. Dashed line represents the FDA recommendation for "high titer" classification ($IC_{50}$~$10^{2.4}$).[3] Details including references, construct type, dose, immunization timeline, route of administration can be found in supplemental **Table S9**.

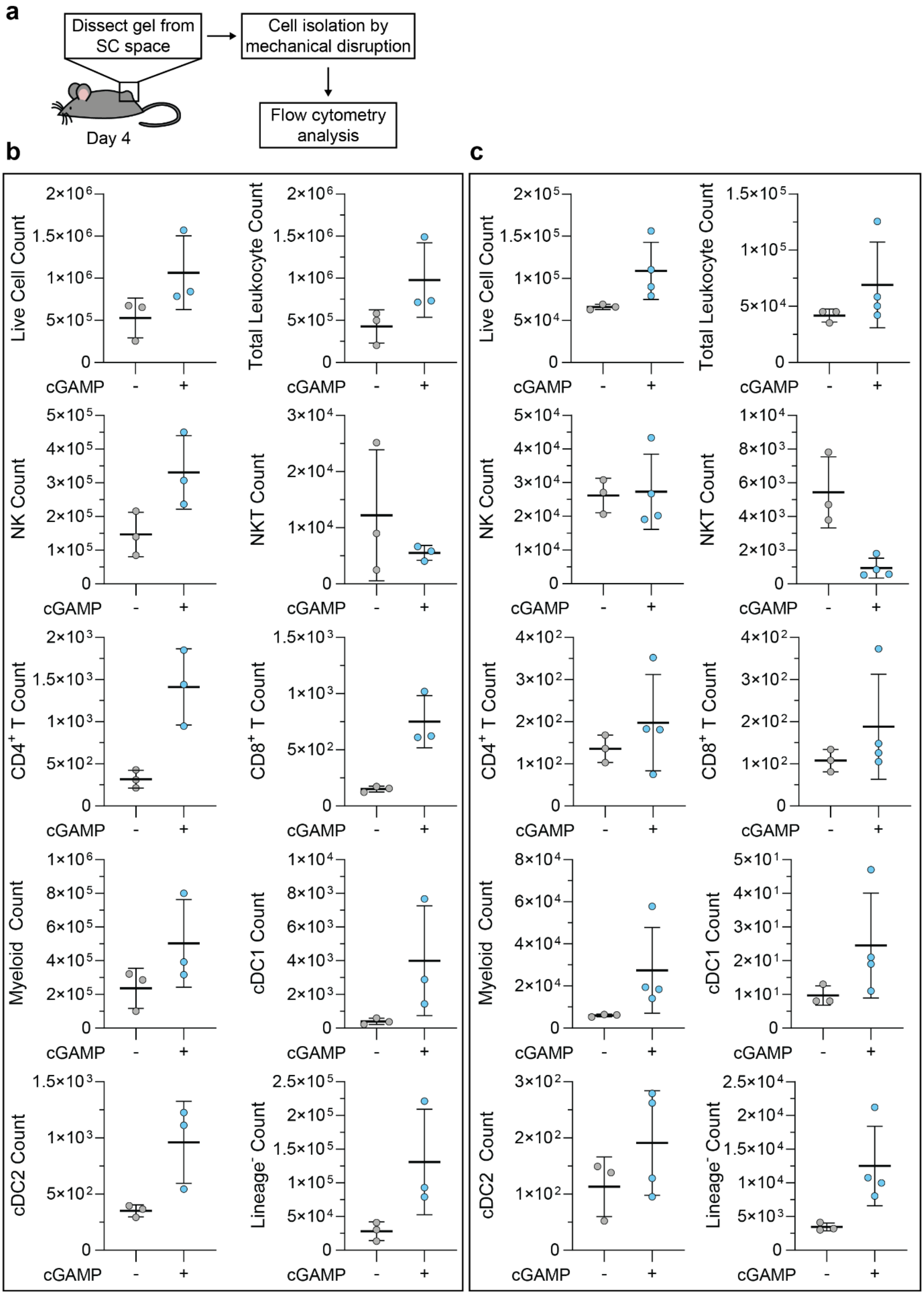


**Figure S4**. Raw characterization of cellular infiltrate to the hydrogel vaccine niche in Figure 4. (A) Schematic of the experimental workflow: four days following injection of hydrogels containing RBD, alum, and the absence or presence of 10 μg cGAMP, gels were excised and dissociated. Total cellular infiltrate was then analyzed by flow cytometer. (B-C) Different experimental rounds of flow cytometry counts of live cells, $CD45^+$ leukocytes, and $CD45^+$ immune subsets. For B, n = 3 mice. For C, n = 3 mice (- cGAMP) and n = 4 mice (+ cGAMP).

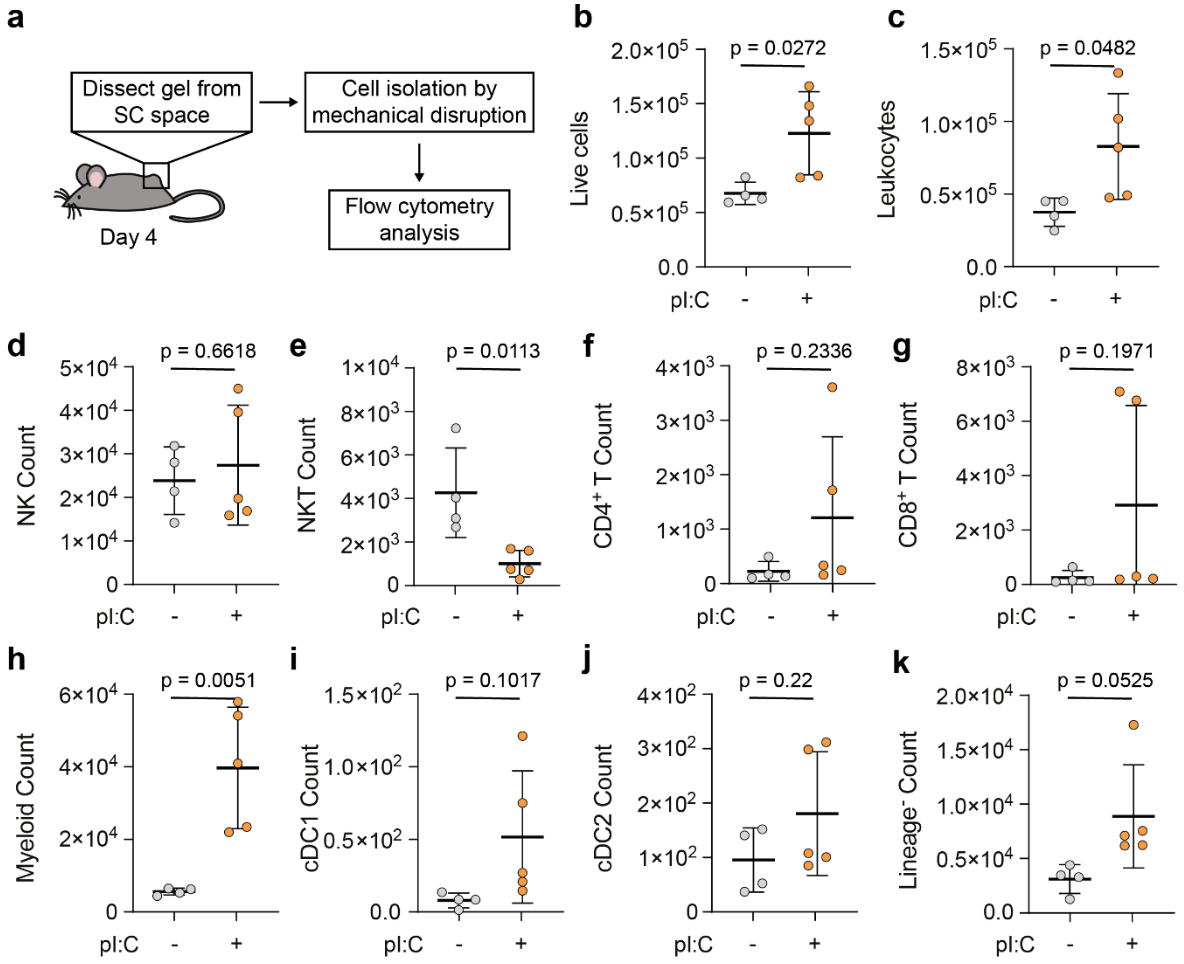


**Figure S5**. Characterization of cellular infiltrate to the hydrogel vaccine niche. (A) Schematic of the experimental workflow: four days following injection of hydrogels containing RBD, alum, and the absence or presence of 50 µg poly(I:C), gels were excised and dissociated. Total cellular infiltrate was then analyzed by flow cytometer. (B-K) Flow cytometry counts of live cells, CD45$^+$ leukocytes, and CD45$^+$ immune subsets. (I) Percentages of major immune subsets among total CD45$^+$ infiltrate, without or with 50 µg poly(I:C). For (B-K), n = 4 mice (- pI:C) and n = 5 mice (+ pI:C) with error bars representing mean ± s.d.; p values are determined by a two-tailed t test.

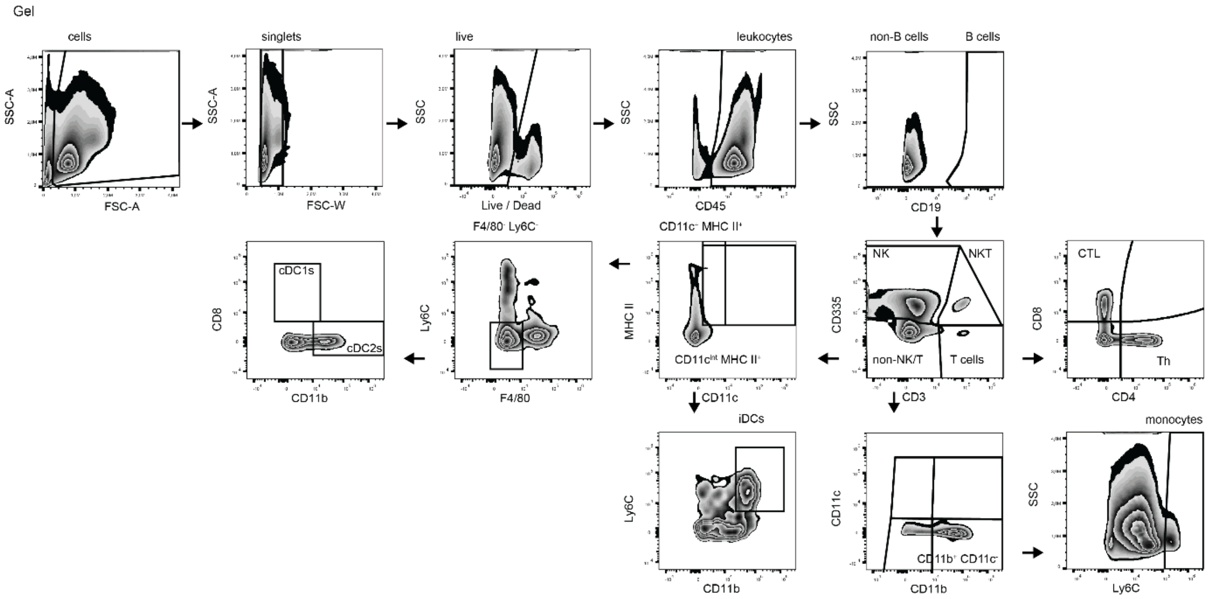

**Figure S6.** Gating strategy for gel samples, in reference to Figure 4.

**Table S1.** P-values from an Ordinary one-way ANOVA with Tukey's multiple comparisons test for 3h IFNα levels (Figures S2b). All groups were compared to each other. ns: p = >0.05; **: p ≤ 0.01

| 3h IFNα levels | Adjusted P Value | significance |
|---|---|---|
| Alum vs Alum + 10 µg cGAMP | 0.9999 | ns |
| Alum vs Alum + 50 µg cGAMP | 0.6089 | ns |
| Alum vs. Gel + Alum + 10 µg cGAMP | 0.9847 | ns |
| Alum vs. Gel + Alum + 50 µg cGAMP | 0.0012 | ** |
| Alum vs. Gel + Alum + 50 µg poly(I:C) | >0.9999 | ns |
| Alum vs Gel + Alum | >0.9999 | ns |
| Alum + 10 µg cGAMP vs Alum + 50 µg cGAMP | 0.8009 | ns |
| Alum + 10 µg cGAMP vs Gel + Alum + 10 µg cGAMP | 0.9992 | ns |
| Alum + 10 µg cGAMP vs Gel + Alum + 50 µg cGAMP | 0.0029 | ** |
| Alum + 10 µg cGAMP vs Gel + Alum + 50 µg poly(I:C) | 0.9999 | ns |
| Alum + 10 µg cGAMP vs Gel + Alum | 0.9999 | ns |
| Alum + 50 µg cGAMP vs Gel + Alum + 10 µg cGAMP | 0.9631 | ns |
| Alum + 50 µg cGAMP vs Gel + Alum + 50 µg cGAMP | 0.0835 | ns |
| Alum + 50 µg cGAMP vs Gel + Alum + 50 µg poly(I:C) | 0.6089 | ns |
| Alum + 50 µg cGAMP vs Gel + Alum | 0.6089 | ns |
| Gel + Alum + 10 µg cGAMP vs Gel + Alum + 50 µg cGAMP | 0.0094 | ** |
| Gel + Alum + 10 µg cGAMP vs Gel + Alum + 50 µg poly(I:C) | 0.9847 | ns |
| Gel + Alum + 10 µg cGAMP vs Gel + Alum | 0.9847 | ns |
| Gel + Alum + 50 µg cGAMP vs Gel + Alum + 50 µg poly(I:C) | 0.0012 | ** |
| Gel + Alum + 50 µg cGAMP vs Gel + Alum | 0.0012 | ** |
| Gel + Alum + 50 µg poly(I:C) vs Gel + Alum | >0.9999 | ns |

**Table S2.** P-values from an Ordinary one-way ANOVA with Tukey's multiple comparisons test for 24h IFNα levels (Figures S2c). All groups were compared to each other. ns: p = >0.05; *: p ≤ 0.05; **: p ≤ 0.01; ***: p ≤ 0.001; ****: p ≤ 0.0001

| 24h IFNα levels | Adjusted P Value | significance |
|---|---|---|
| Alum vs Alum + 10 µg cGAMP | 0.7037 | ns |
| Alum vs Alum + 50 µg cGAMP | 0.0045 | ** |
| Alum vs. Gel + Alum + 10 µg cGAMP | 0.9697 | ns |
| Alum vs. Gel + Alum + 50 µg cGAMP | 0.0002 | *** |
| Alum vs. Gel + Alum + 50 µg poly(I:C) | 0.9916 | ns |
| Alum vs Gel + Alum | >0.9999 | ns |
| Alum + 10 µg cGAMP vs Alum + 50 µg cGAMP | 0.2301 | ns |
| Alum + 10 µg cGAMP vs Gel + Alum + 10 µg cGAMP | 0.9705 | ns |
| Alum + 10 µg cGAMP vs Gel + Alum + 50 µg cGAMP | 0.0495 | * |
| Alum + 10 µg cGAMP vs Gel + Alum + 50 µg poly(I:C) | 0.9244 | ns |
| Alum + 10 µg cGAMP vs Gel + Alum | 0.5528 | ns |
| Alum + 50 µg cGAMP vs Gel + Alum + 10 µg cGAMP | 0.0108 | * |
| Alum + 50 µg cGAMP vs Gel + Alum + 50 µg cGAMP | 0.9999 | ns |
| Alum + 50 µg cGAMP vs Gel + Alum + 50 µg poly(I:C) | 0.0061 | ** |
| Alum + 50 µg cGAMP vs Gel + Alum | 0.0007 | *** |
| Gel + Alum + 10 µg cGAMP vs Gel + Alum + 50 µg cGAMP | 0.0002 | *** |
| Gel + Alum + 10 µg cGAMP vs Gel + Alum + 50 µg poly(I:C) | >0.9999 | ns |
| Gel + Alum + 10 µg cGAMP vs Gel + Alum | 0.9240 | ns |
| Gel + Alum + 50 µg cGAMP vs Gel + Alum + 50 µg poly(I:C) | 0.0001 | *** |
| Gel + Alum + 50 µg cGAMP vs Gel + Alum | <0.0001 | **** |
| Gel + Alum + 50 µg poly(I:C) vs Gel + Alum | 0.9770 | ns |

**Table S3.** P-values from a 2-way ANOVA with Dunnett's multiple comparisons test for specific IgG titer time points where Gel + Alum + 10 µg cGAMP, Gel + Alum + 50 µg cGAMP, and Gel + Alum + 50 µg poly(I:C) groups are compared to the Gel + Alum control. ns: p = >0.05; *: p ≤ 0.05; **: p ≤ 0.01; ***: p ≤ 0.001

| Week 2<br>IgG titers [$\log_{10}$][a)] | Adjusted P Value | significance |
|---|---|---|
| Gel + Alum vs. Gel + Alum + 10 µg cGAMP | 0.0004 | *** |
| Gel + Alum vs. Gel + Alum + 50 µg cGAMP | 0.0010 | *** |
| Gel + Alum vs. Gel + Alum + 50 µg poly(I:C) | 0.3424 | ns |

| Week 4<br>IgG titers [$\log_{10}$][a)] | Adjusted P Value | significance |
|---|---|---|
| Gel + Alum vs. Gel + Alum + 10 µg cGAMP | 0.0358 | * |
| Gel + Alum vs. Gel + Alum + 50 µg cGAMP | 0.0463 | * |
| Gel + Alum vs. Gel + Alum + 50 µg poly(I:C) | 0.5083 | ns |

| Week 8<br>IgG titers [$\log_{10}$][a)] | Adjusted P Value | significance |
|---|---|---|
| Gel + Alum vs. Gel + Alum + 10 µg cGAMP | 0.0468 | * |
| Gel + Alum vs. Gel + Alum + 50 µg cGAMP | 0.1446 | ns |
| Gel + Alum vs. Gel + Alum + 50 µg poly(I:C) | 0.2592 | ns |

| Week 12<br>IgG titers [$\log_{10}$][a)] | Adjusted P Value | significance |
|---|---|---|
| Gel + Alum vs. Gel + Alum + 10 µg cGAMP | 0.0306 | * |
| Gel + Alum vs. Gel + Alum + 50 µg cGAMP | 0.0015 | ** |
| Gel + Alum vs. Gel + Alum + 50 µg poly(I:C) | >0.9999 | ns |

[a)] Statistical analyses are run on logged titer values

**Table S4.** P-values from a 2-way ANOVA with Tukey's multiple comparisons test for specific IgG titer time points where all combinations of Alum + 10 µg cGAMP, Alum + 50 µg cGAMP, Gel + Alum + 10 µg cGAMP, Gel + Alum + 50 µg cGAMP, and Gel + Alum + 50 µg Poly(I:C) were compared. Only relevant comparisons (matched dose or delivery system) are shown in the table below. ns: p = >0.05; *: p ≤ 0.05; **: p ≤ 0.01; ****: p ≤ 0.0001

| Week 2<br>IgG titers [$\log_{10}$][a)] | Adjusted P Value | significance |
|---|---|---|
| Alum + 10 µg cGAMP vs. Alum + 50 µg cGAMP | >0.9999 | ns |
| Alum + 10 µg cGAMP vs. Gel + Alum + 10 µg cGAMP | 0.9552 | ns |
| Alum + 50 µg cGAMP vs. Gel + Alum + 50 µg cGAMP | 0.0087 | ** |
| Gel + Alum + 10 µg cGAMP vs. Gel + Alum + 50 µg cGAMP | 0.0242 | * |
| Gel + Alum + 50 µg cGAMP vs. Gel + Alum + 50 µg Poly(I:C) | 0.1572 | ns |

| Week 4<br>IgG titers [$\log_{10}$][a)] | Adjusted P Value | significance |
|---|---|---|
| Alum + 10 µg cGAMP vs. Alum + 50 µg cGAMP | >0.9999 | ns |
| Alum + 10 µg cGAMP vs. Gel + Alum + 10 µg cGAMP | <0.0001 | **** |
| Alum + 50 µg cGAMP vs. Gel + Alum + 50 µg cGAMP | <0.0001 | **** |
| Gel + Alum + 10 µg cGAMP vs. Gel + Alum + 50 µg cGAMP | 0.9993 | ns |
| Gel + Alum + 50 µg cGAMP vs. Gel + Alum + 50 µg Poly(I:C) | 0.7212 | ns |

| Week 8<br>IgG titers [$\log_{10}$][a)] | Adjusted P Value | significance |
|---|---|---|
| Alum + 10 µg cGAMP vs. Alum + 50 µg cGAMP | 0.6551 | ns |
| Alum + 10 µg cGAMP vs. Gel + Alum + 10 µg cGAMP | <0.0001 | **** |
| Alum + 50 µg cGAMP vs. Gel + Alum + 50 µg cGAMP | <0.0001 | **** |
| Gel + Alum + 10 µg cGAMP vs. Gel + Alum + 50 µg cGAMP | 0.9277 | ns |
| Gel + Alum + 50 µg cGAMP vs. Gel + Alum + 50 µg Poly(I:C) | 0.9980 | ns |

| Week 12<br>IgG titers [$\log_{10}$][a)] | Adjusted P Value | significance |
|---|---|---|
| Alum + 10 µg cGAMP vs. Alum + 50 µg cGAMP | 0.0566 | ns |
| Alum + 10 µg cGAMP vs. Gel + Alum + 10 µg cGAMP | <0.0001 | **** |
| Alum + 50 µg cGAMP vs. Gel + Alum + 50 µg cGAMP | <0.0001 | **** |
| Gel + Alum + 10 µg cGAMP vs. Gel + Alum + 50 µg cGAMP | 0.6111 | ns |
| Gel + Alum + 50 µg cGAMP vs. Gel + Alum + 50 µg Poly(I:C) | 0.0046 | ** |

[a)]Statistical analyses are run on logged titer values

**Table S5.** P-values from a 2- way ANOVA with Dunnett's multiple comparisons test for specific IgG titer time points where Alum + 10 µg cGAMP and Alum + 50 µg cGAMP groups are compared to the Alum control. ns: p = >0.05; ***: $p \leq 0.001$

| Week 2<br>IgG titers [$\log_{10}$][a)] | Adjusted P Value | significance |
|---|---|---|
| Alum vs. Alum + 10 µg cGAMP | 0.7345 | ns |
| Alum vs. Alum + 50 µg cGAMP | 0.7345 | ns |

| Week 4<br>IgG titers [$\log_{10}$][a)] | Adjusted P Value | significance |
|---|---|---|
| Alum vs. Alum + 10 µg cGAMP | 0.5948 | ns |
| Alum vs. Alum + 50 µg cGAMP | 0.5948 | ns |

| Week 8<br>IgG titers [$\log_{10}$][a)] | Adjusted P Value | significance |
|---|---|---|
| Alum vs. Alum + 10 µg cGAMP | 0.4804 | ns |
| Alum vs. Alum + 50 µg cGAMP | 0.9769 | ns |

| Week 12<br>IgG titers [$\log_{10}$][a)] | Adjusted P Value | significance |
|---|---|---|
| Alum vs. Alum + 10 µg cGAMP | 0.3080 | ns |
| Alum vs. Alum + 50 µg cGAMP | 0.0003 | *** |

[a)]Statistical analyses are run on logged titer values

**Table S6.** P-values from a 2-way ANOVA with Dunnett's multiple comparisons test for post-boost (week 12) IgG1 and IgG2c titers where Gel + Alum + 10 µg cGAMP, Gel + Alum + 50 µg cGAMP, and Gel + Alum + 50 µg poly(I:C) groups are compared to the Gel + Alum control. ns: p = >0.05; *: p ≤ 0.05; **: p ≤ 0.01; ***: p ≤ 0.001; ****: p ≤ 0.0001

| Week 12<br>IgG1 titers [$log_{10}$][a)] | Adjusted P Value | significance |
|---|---|---|
| Gel + Alum vs. Gel + Alum + 10 µg cGAMP | 0.0006 | *** |
| Gel + Alum vs. Gel + Alum + 50 µg cGAMP | <0.0001 | **** |
| Gel + Alum vs. Gel + Alum + 50 µg poly(I:C) | >0.9999 | ns |

| Week 12<br>IgG2c titers [$log_{10}$][a)] | Adjusted P Value | significance |
|---|---|---|
| Gel + Alum vs. Gel + Alum + 10 µg cGAMP | 0.0354 | * |
| Gel + Alum vs. Gel + Alum + 50 µg cGAMP | 0.0002 | *** |
| Gel + Alum vs. Gel + Alum + 50 µg poly(I:C) | 0.9689 | ns |

[a)]Statistical analyses are run on logged titer values

**Table S7.** P-values from a 2-way ANOVA with Dunnett's multiple comparisons test for post-boost (week 12) IgG titers where Alpha (B.1.1.7, UK), Beta (B.1.351, South Africa), and Delta (B.1.617.2) spike variants are compared to wildtype spike for a given treatment group. ns: p = >0.05; *: p ≤ 0.05; **: p ≤ 0.01

| Week 12<br>IgG titers [$log_{10}$][a)] | Adjusted P Value | significance |
|---|---|---|
| Alum: WT spike vs. Alpha variant | 0.8555 | ns |
| Alum: WT spike vs. Beta variant | 0.0452 | * |
| Alum: WT spike vs. Delta variant | 0.9997 | ns |
| Alum + 10 µg cGAMP: WT spike vs. Alpha variant | 0.7794 | ns |
| Alum + 10 µg cGAMP: WT spike vs. Beta variant | 0.0089 | ** |
| Alum + 10 µg cGAMP: WT spike vs. Delta variant | 0.9505 | ns |
| Alum + 50 µg cGAMP: WT spike vs. Alpha variant | 0.8724 | ns |
| Alum + 50 µg cGAMP: WT spike vs. Beta variant | 0.0170 | * |
| Alum + 50 µg cGAMP: WT spike vs. Delta variant | 0.9980 | ns |
| Gel + Alum: WT spike vs. Alpha variant | 0.6638 | ns |
| Gel + Alum: WT spike vs. Beta variant | 0.0271 | * |
| Gel + Alum: WT spike vs. Delta variant | 0.9980 | ns |
| Gel + Alum + 10 µg cGAMP: WT spike vs. Alpha variant | 0.7336 | ns |
| Gel + Alum + 10 µg cGAMP: WT spike vs. Beta variant | 0.0414 | * |
| Gel + Alum + 10 µg cGAMP: WT spike vs. Delta variant | 0.9994 | ns |
| Gel + Alum + 50 µg cGAMP: WT spike vs. Alpha variant | 0.5557 | ns |
| Gel + Alum + 50 µg cGAMP: WT spike vs. Beta variant | 0.0120 | * |
| Gel + Alum + 50 µg cGAMP: WT spike vs. Delta variant | >0.9999 | ns |
| Gel + Alum + 50 µg poly(I:C): WT spike vs. Alpha variant | 0.7816 | ns |
| Gel + Alum + 50 µg poly(I:C): WT spike vs. Beta variant | 0.0190 | * |
| Gel + Alum + 50 µg poly(I:C): WT spike vs. Delta variant | 0.8873 | ns |

[a)]Statistical analyses are run on logged titer values

**Table S8.** P-values from an Ordinary one-way ANOVA with Dunnett's multiple comparisons test for post-boost (week 12) serum $IC_{50}$ values from a reporter lentivirus pseuodotyped SARS-CoV-2 spike neutralization assay. Gel + Alum + 10 µg cGAMP, Gel + Alum + 50 µg cGAMP, and Gel + Alum + 50 µg poly(I:C) groups are compared to the Gel + Alum control. ns: p = >0.05.

| Construct | Immunization timeline | Dose | Neutralization assay | Route of administration | Adjuvant |
|---|---|---|---|---|---|
| WT RBD in cGAMP gel | Day 0 & 56 | 10 µg | Pseudovirus | Subcutaneous | cGAMP |
| Rpk9-I53-50[4] | Day 0 & 21 | 0.9 µg | Pseudovirus | Intramuscular | AddaVax |
| RBD-L452K-F490W[5] | Day 0 & 21 | 5 µg | Pseudovirus | Subcutaneous | CpG & Alum |
| SΔFer[2] | Day 0 & 21 | 10 µg | Pseudovirus | Subcutaneous | Quil-A & MPLA |
| RBD-16GS-I53-50[6] | Day 0 & 21 | 5 µg | Pseudovirus | Intramuscular | AddaVax |
| CpG + Alum + Gel[1] | Day 0 & 56 | 20 µg | Pseudovirus | Subcutaneous | CpG ODN 1826 |
| BNT162b1[7] | Day 0 | 5 µg mRNA | Pseudovirus | Intramuscular | None |
| RBDsurf[8] | Day 0 & 21 | 10 µg | Viral | Subcutaneous | MPLA PS |
| BNT162b2[7] | Day 0 | 5 µg mRNA | Pseudovirus | Intramuscular | None |

**Table S9.** References, construct type, timeline, dose, neutralization assay type, route of administration, and adjuvant(s) for all immunizations shown in **Figure S3**. **Table S8.**

| Week 12<br>$IC_{50}$ [reciprocal serum dilution] | Adjusted P Value | significance |
|---|---|---|
| Gel + Alum vs. Gel + Alum + 10 µg cGAMP | 0.2405 | ns |
| Gel + Alum vs. Gel + Alum + 50 µg cGAMP | 0.0980 | ns |
| Gel + Alum vs. Gel + Alum + 50 µg poly(I:C) | >0.9999 | ns |

**Table S10.** Information on flow cytometry antibodies.

| **Antigen** | **Manufacturer** | **Clone** | **Fluorophore** | **Dilution** |
|---|---|---|---|---|
| CD11c | BD Biosciences | HL3 | BUV395 | 1:200 |
| CD4 | BD Biosciences | GK1.5 | BUV805 | 1:200 |
| CD11b | BioLegend | M1/70 | BV421 | 1:400 |
| F4/80 | BioLegend | BM8 | BV510 | 1:400 |
| Ly-6C | BioLegend | HK1.4 | BV570 | 1:200 |
| CD335 | BioLegend | 29A1.4 | BV605 | 1:800 |
| CD8α | BioLegend | 53-6.7 | BV785 | 1:200 |
| pSTING | CST | D1C4T | | 1:200 |
| I-A/I-E | BioLegend | M5/114.15.2 | PerCP-Cy5.5 | 1:100 |
| CD3ε | eBioscience | eBio500A2 | PerCP-eF710 | 1:200 |
| His6 | BioLegend | J095G46 | PE | 1:200 |
| CD19 | BioLegend | 6D5 | PE-Cy7 | 1:200 |
| pIRF3 | CST | D6O1M | AF647 | 1:50 |
| CD45 | BioLegend | 30-F11 | AF700 | 1:800 |
